# Critical Sequence Hot-spots for Binding of nCOV-2019 to ACE2 as Evaluated by Molecular Simulations

**DOI:** 10.1101/2020.06.27.175448

**Authors:** Mahdi Ghorbani, Bernard R. Brooks, Jeffery B. Klauda

**Affiliations:** Department of Chemical and Biomolecular Engineering, University of Maryland, College Park, MD 20742, USA; Biophysics Graduate Program, University of Maryland, College Park, MD 20742, USA; Laboratory of Computational Biology, National, Heart, Lung and Blood Institute, National Institutes of Health, Bethesda, Maryland 20824, USA

## Abstract

The novel coronavirus (nCOV-2019) outbreak has put the world on edge, causing millions of cases and hundreds of thousands of deaths all around the world, as of June 2020, let alone the societal and economic impacts of the crisis. The spike protein of nCOV-2019 resides on the virion’s surface mediating coronavirus entry into host cells by binding its receptor binding domain (RBD) to the host cell surface receptor protein, angiotensin converter enzyme (ACE2). Our goal is to provide a detailed structural mechanism of how nCOV-2019 recognizes and establishes contacts with ACE2 and its difference with an earlier coronavirus SARS-COV in 2002 via extensive molecular dynamics (MD) simulations. Numerous mutations have been identified in the RBD of nCOV-2019 strains isolated from humans in different parts of the world. In this study, we investigated the effect of these mutations as well as other Ala-scanning mutations on the stability of RBD/ACE2 complex. It is found that most of the naturally-occurring mutations to the RBD either strengthen or have the same binding affinity to ACE2 as the wild-type nCOV-2019. This may have implications for high human-to-human transmission of coronavirus in regions where these mutations have been found as well as any vaccine design endeavors since these mutations could act as antibody escape mutants. Furthermore, in-silico Ala-scanning and long-timescale MD simulations, highlight the crucial role of the residues at the interface of RBD and ACE2 that may be used as potential pharmacophores for any drug development endeavors. From an evolutional perspective, this study also identifies how the virus has evolved from its predecessor SARS-COV and how it could further evolve to become more infectious.

## Introduction

The novel coronavirus (nCOV-2019) outbreak emerging from China has become a global pandemic and a major threat for human public health. According to World Health Organization (WHO) as of June 27^th^ 2020, there has been about 10 million confirmed cases and approaching 500,000 deaths due to coronavirus in the world.^1–2^ Much of the human population including the United States of America were under lockdown or official stay-at-home orders to minimize the continued spread of the virus.

Coronaviruses are a family of single-stranded enveloped RNA viruses. Phylogenetic analysis of coronavirus genome has shown that nCOV-2019 belongs to the beta-coronavirus family, which also includes MERS-COV, SARS-COV and bat-SARS-related coronaviruses.^3–4^ It is worth mentioning that SARS-COV, which was widespread in 2002 caused more than 8,000 cases and about 800 deaths and MERS-COV (middle east respiratory syndrome coronavirus) in 2012 also spread in more than 25 countries, causing about 2,500 cases and more than 850 deaths. (www.who.int/health-topics/coronavirus).^5^

In all coronaviruses, a homotrimeric spike glycoprotein on the virion’s envelope mediates coronavirus entry into host cells through a mechanism of receptor binding followed by fusion of viral and host membranes.^3,6^ Coronavirus spike protein contains two functional subunits S1 and S2. The S1 subunit is responsible for binding to host cell receptor, and the S2 subunit is responsible for fusion of viral and host cell membranes.^3,7^ The spike protein in nCOV-2019 exists in a meta-stable pre-fusion conformation that undergoes a substantial conformational rearrangement to fuse the viral membrane with the host cell membrane.^7,8^ nCOV-2019 is closely related to bat coronavirus RaTG13 with about 93.1% sequence similarity in the spike protein gene. The sequence similarity of nCOV-2019 and SARS-COV is less than 80% in the spike genome.^2^ S1 subunit in the spike protein includes a receptor binding domain (RBD) that recognizes and binds to the host cells receptor. The RBD of nCOV-2019 shares 72.8% sequence identity to SARS-COV RBD and the root mean squared deviation (RMSD) for the structure between the two proteins is 1.2 *Å*, which shows the high structural similarity.^4,8,9^ Experimental binding affinity measurements using Surface Plasmon Resonance (SPR) have shown that nCOV-2019 spike protein binds its receptor human angiotensin converter enzyme (ACE2) with 10 to 20 fold higher affinity than SARS-COV binding to ACE2.^7^ Based on the sequence similarity between RBD of nCOV-2019 and SARS-COV and also the tight binding between RBD of nCOV-2019 and ACE2, it is most probable that nCOV-2019 uses this receptor on human cells to gain entry into the body.^3,6,7,10^

The spike protein and specifically the RBD domain in coronaviruses have been a major target for therapeutic antibodies. However, no monoclonal antibodies targeted to RBD have been able to bind efficiently and neutralize nCOV-2019.^7,11^ The core of nCOV-2019 RBD is a 5-stranded antiparallel *β*-sheet with connected short α-helices and loops (Figure 1). Binding interface of nCOV-2019 and SARS-COV with ACE2 are very similar with less than 1.3 Å RMSD. An extended insertion inside the core containing short strands, α-helices and loops called the receptor binding motif (RBM) makes all the contacts with ACE2. In nCOV-2019 RBD, the RBM forms a concave surface with a ridge loop on one side and it binds to a convex exposed surface of ACE2. The overlay of SARS and nCOV-2019 RBD proteins are shown in Figure 1A. The binding interface in nCOV-2019 contains loops *L*_1_ to *L*_4_ and short *β*-strands *β*_5_ and *β*_6_ and a short helix *α*_5_. The location of RBM in nCOV-2019 RBD as well as different helices, strands and loops is shown in Figure 1B.

**Figure 1.**
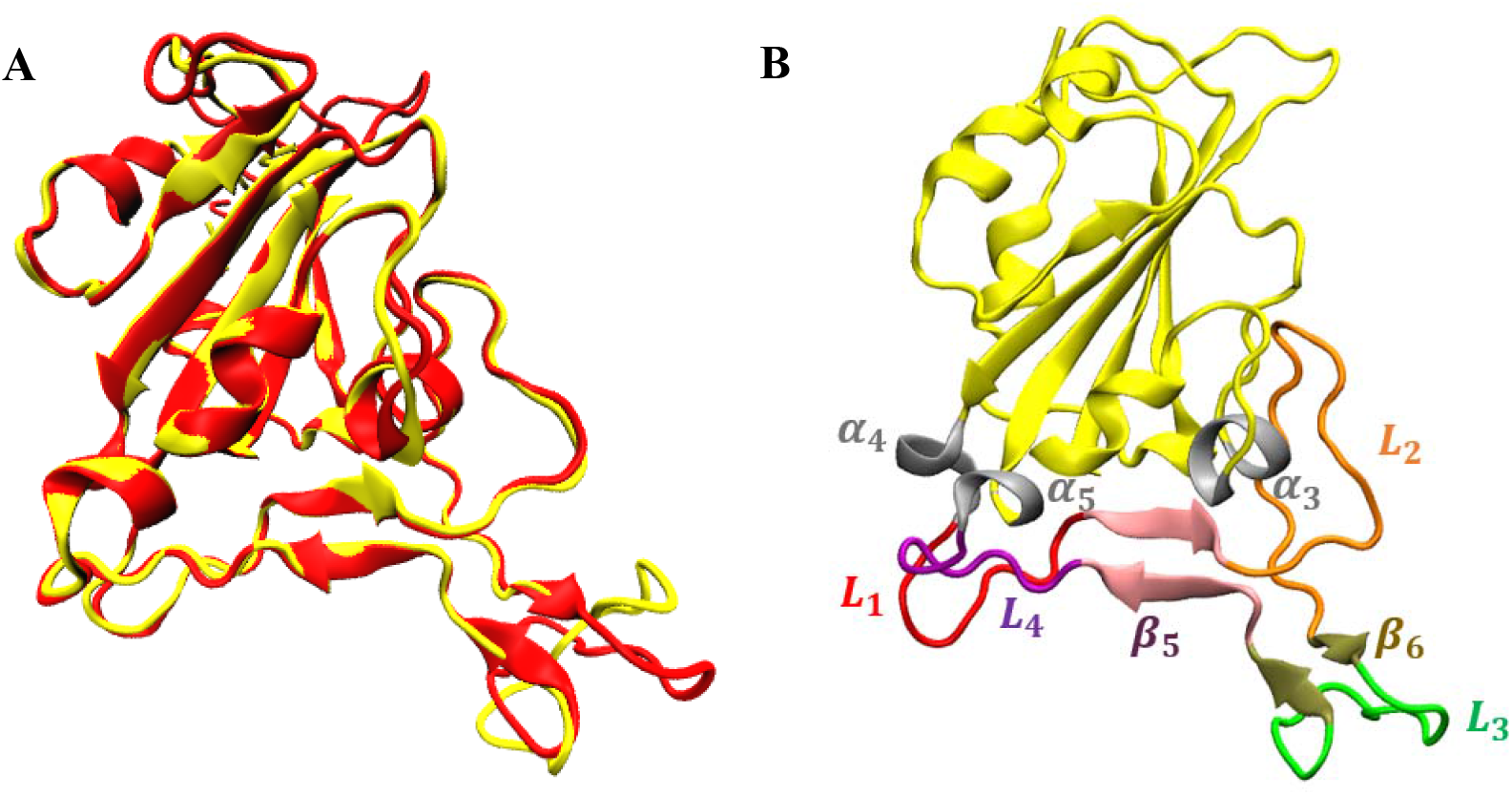
A) superposition of the RBD of SARS-COV (yellow) and nCOV-2019(red). B) different regions in the binding domain of nCOV-2019 defining the extended loop (non-yellow).

The sequence alignment between SARS-COV in human, SARS civet, Bat RaTG13 coronavirus and nCOV-2019 in the RBM is shown in Figure 2. There is a 50% sequence similarity between the RBM of nCOV-2019 and SARS-COV. RBM mutations played an important role in the SARS epidemic in 2002.^12,3^ Two mutations in the RBM of SARS-2002 from SARS-Civet were observed from strains of these viruses. These two mutations were K479N and S487T. These two residues are close to the virus binding hotspots in ACE2 including hotspot-31 and hotspot-353. Hotspot-31 centers on the salt-bridge between K31-E35 and hotspot-353 is centered on the salt-bridge between K353-E358 on ACE2. Residues K479 and S487 in SARS-Civet are in close proximity with these hotspots and mutations at these residues caused SARS to bind ACE2 with significantly higher affinity than SARS-Civet and played a major role in civet-to-human and human-to-human transmission of SARS coronavirus in 2002.^3,13–15^ Numerous mutations in the interface of SARS-COV RBD and ACE2 from different strains of SARS isolated from humans in 2002 have been identified and the effect of these mutations on binding ACE2 have been investigated by surface plasmon resonance.^16,17^ Two identified RBD mutations (Y442F and L472F) increased the binding affinity of SARS-COV to ACE2 and two mutations (N479K, T487S) decreased the binding affinity. It was demonstrated that these mutations were viral adaptations to either human or civet ACE2.^16,17^ A pseudotyped viral infection assay of the interaction between different spike proteins and ACE2 confirmed the correlation between high affinity mutants and their high infection.^16^ Further investigation of RBD residues in binding of SARS-COV and ACE2 was performed through ala-scanning mutagenesis, which resulted in identification of 10 residues that reduce binding affinity to ACE2 upon mutation to alanine. These residues are K390, R429, D429, D454, I455, N473, F483, Q492, Y494 and R495.^18^ RBD mutations have also been identified in MERS-COV, which affected their affinity to receptor (DPP4) on human cells.^17^ Multiple monoclonal antibodies have been developed for SARS since 2002 that neutralized spike glycoprotein on SARS-COV surface.^19–23^ However, multiple escape mutations exist in the RBD of SARS-COV that affect neutralization with antibodies, which led to the use of a cocktail of antibodies as robust treatment.^24^

**Figure 2.**
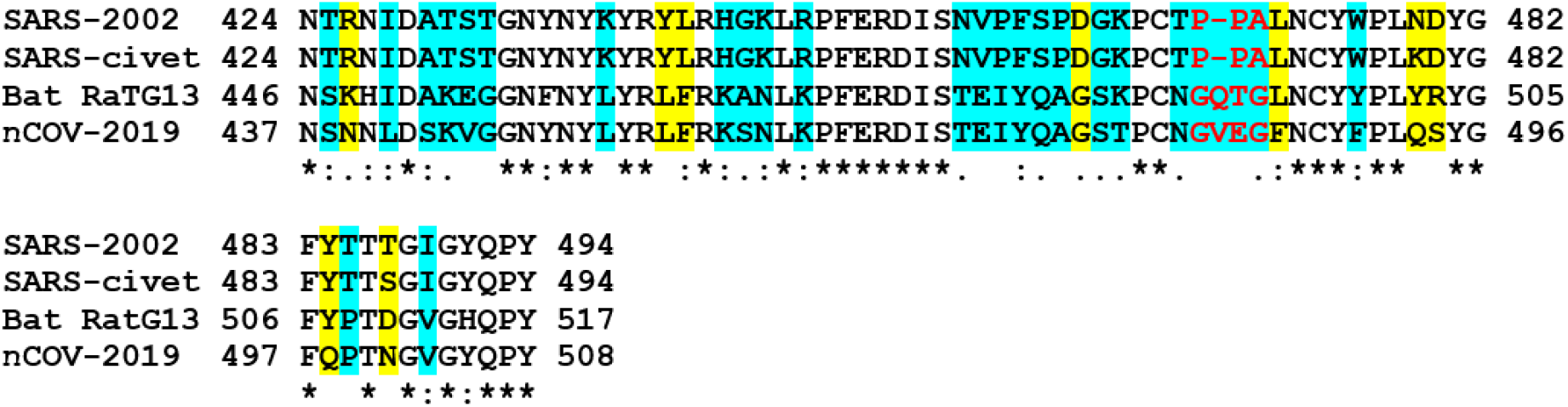
Sequence comparison of the receptor binding motif (RBM) in SARS-2002, SARS-civet, Bat RaTG13 and nCOV-2019. The mutations from SARS-2002 to nCOV-2019 are marked with blue. Important mutations in RBM are marked with yellow. Red color shows the 3-resdiue motif in SARS and Civet and 4-residue motif in RaTG13 and nCOV-2019.

Full genome analysis of nCOV-2019 in different countries and the receptor binding surveillance has shown multiple mutations in the RBD of glycosylated spike. The GISAID database^25^ (www.gisaid.org/) contains genomes on nCOV-2019 from researchers across the world since December 2019. Latest report by the GISAID database on June 2020 have shown 25 different variants of RBD from strains of nCOV-2019 collected from different countries along with the number of occurrences in these regions which is listed below for the seven most occurring mutations:

> 213x **N439K** (211 Scotland, England, Romania), 65x **T478I** in England, 30x **V483A** (26 USA/WA, 2 USA/UN, USA/CT, England), 10x **G476S** (8 USA/WA, USA/OR, Belgium), 7x **S494P** (3 USA/MI, England, Spain, India, Sweden), 5x **V483F** (4x Spain, England), 4x **A475V** (2 USA/AZ, USA/NY, Australia/NSW.

It is not known whether these mutations are linked to the severity of coronavirus in these regions. The focus of this article is to elucidate the differences between the interface of SARS-COV and nCOV-2019 with ACE2 to understand with atomic resolution the interaction mechanism and hotspot residues at the RBD/ACE2 interface using long-timescale molecular dynamics (MD) simulation. An alanine-scanning mutagenesis in the RBM of nCOV-2019 helped to identify the key residues in the interaction, which could be used as potential pharmacophores for future drug development. Furthermore, we performed molecular simulations on the seven most common mutations found from surveillance of RBD mutations N439K, T478I, V483A,G476S, S494P, V483F and A475V. From an evolutionary perspective this study shows the residues in which the virus might further evolve to be even more dangerous to human health.

## Methods

### Sequence comparison and mutant preparation

nCOV-2019 shares 76% sequence similarity with SARS-2002 spike protein, 73% sequence identity for RBD and 50% for the RBM.^1^ Bat coronavirus RaTG13 seems to be the closest relative of nCOV-2019 sharing about 93% sequence identity in the spike protein.^6^ The sequence alignment of SARS-2002 (accession number: AFR58742), SARS-civet (accession number: AY304486.1), Bat RaTG13 (accession number: MN996532.1) and nCOV-2019 (accession number: MN908947.1) are in Figure 2.^26^ To investigate the roles of critical mutations on the complex stability of nCOV-2019 with ACE2, mCSM-PPI2 webserver^27^ was used to the residues in nCOV-2019 that are at the interface with ACE2. This method uses a graph-based signature framework and predicts the effect of alanine substitution at interface residues on the binding energy of the complex. *21* different residues were identified to be in contact with ACE2 and were chosen to do further MD simulation. On the other hand, mutations are also observed in RBD domain from full genome analysis of different nCOV-2019 variants collected from different countries compiled in GISAID database.^25^ The mutations selected are listed in Table S1 along with their location in RBD.

### Molecular dynamics simulations

The crystal structure of nCOV-2019 in complex with hACE2 (pdb id:6M0J)^28^ as well as SARS-COV complex with human ACE2 (pdb id: 6ACJ)^29^ were obtained from RCSB (www.rcsb.org).^30^ The RBD domain of nCOV-2019 comprises of 194 residues (333-526) and SARS-COV includes 190 residues (323-512). ACE2 protein contains 597 residues (19-615) in the complex structure for both systems. All the structures including nCOV-2019, SARS-COV and all 21 alanine substitutions of nCOV-2019 were prepared and solvated with the CHARMM-GUI webserver.^31^ A TIP3P water model was used for the solvent and Param99SB-ILDN AMBER force field (FF)^32^was used for all the complexes. POT ions were added to all systems to neutralize the charges on the proteins. Each system contained about 260,000 atoms.

5000 steps of energy minimizations were done using the steepest descent algorithm. In all steps the LINCS algorithm was used to constraint all bonds containing hydrogen atoms and a time step of 2 fs was used as the integration time step.^33^ Equilibration of all systems were performed in three steps. In the first step, 100,000 steps of simulation were performed using a velocity-rescaling thermostat to maintain the temperature at 310K with a 0.1 ps coupling constant in NVT ensemble under periodic boundary conditions and harmonic restraints on the backbone and sidechain atoms of the complex.^34^ The velocity rescaling thermostat was used in all other steps of simulation. In the next step, we performed 300,000 steps in the isothermal-isobaric NPT ensemble at temperature of 310K and pressure of 1bar using a Berendsen barostat.^35^ This was done by decreasing the force constant of the restraint on backbone and side chain atoms of the complex from 1000 to 100 and finally to 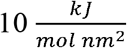. Berendsen barostat was only used for the equilibration step due to usefulness in rapidly correcting density. In the next step the restraints were removed, and the systems were subjected to 1,000,000 steps of MD simulation under NPT ensemble.

In the production run, harmonic restraints were removed and all the systems were simulated using a NPT ensemble where the pressure was maintained at 1bar using the Parrinello-Rahman barostat^36^ with a compressibility of 4.5 × 10^-5^ bar^-1^ and a coupling constant of 0.5ps. It is important to note that all the Berendsen barostat was only used for the equilibration step as it was shown that this barostat can cause unrealistic temperature gradients.^37^ The production run lasted for 500ns for SARS-COV and nCOV-2019 complexes and 300ns for all the mutants using with a 2fs timestep and the particle-mesh Ewald (PME)^36^ for long range electrostatic interactions using GROMACS 2018.3 package.^38^ All mutant systems were constructed as described before and all complexes ran for 300ns of production run.

#### Gibbs free energy and correlated motions

The last 400ns of simulation was used to explore the dominant motions in SARS-COV, nCOV-2019 and last 200ns for all its mutants using principal component analysis (PCA) as part of the quasiharmonic analysis method.^39^ For this method the rotational and translational motions of RBD of all systems were eliminated by fitting to a reference (crystal) structure. Next, 4,000 snapshots from the last 400ns of SARS-COV and nCOV-2019 systems and 2,000 snapshots from last 200ns of all mutant systems were taken to generate the covariance matrix between *C_α_* atoms of RBD. Diagonalization of this matrix resulted in a diagonal matrix of eigenvalues and their corresponding eigenvectors.^38,40^ The first eigenvector which indicate the first principal component was used to visualize the dominant global motions of all complexes through porcupine plots using the (PorcupinePlot.tcl) script in vmd.

The principal components were used to calculate and plot the approximate free energy landscape (aFEL). We refer to the free energy landscape produced by this approach to be approximate in that the ensemble with respect to the first few PC’s (lowest frequency quasiharmonic modes) is not close to convergence, but the analysis can still provide valuable information and insight. g_sham, g_covar and g_anaeig functions in GROMACS^41^ were used to obtain principal components and aFEL. In each aFEL the deep valleys represent the most stable conformations separated by some intermediate states.

The dynamic cross-correlation maps (DCCM) were obtained using MD_TASK package to identify the correlated motions of RBD residues.^42^ In DCCM the cross-correlation matrix *C_ij_* is obtained from displacement of backbone *C_α_* atoms in a time interval △t. The DCCM was constructed using the last 400ns of SARS-COV and nCOV-2019 systems and last 200ns of all mutant systems with a 100ps time interval.

#### Binding free energy from MMPBSA method

The Molecular Mechanics Poisson-Boltzmann Surface Area (MMPBSA) method was applied to calculate the binding energy between RBD and ACE2 in all complexes.^43,44^ for SARS-COV and nCOV-2019 system, 200 snapshots of the last 400ns and for all other systems, 100 snapshots of the last 200ns simulation were used to for the calculation of binding free energies with an interval of 2ns. The binding free energy of a ligand-receptor complex can be calculated as:

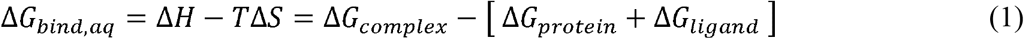

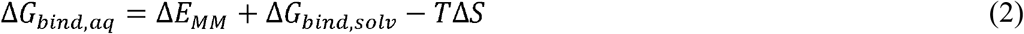

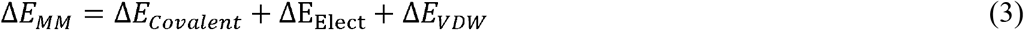

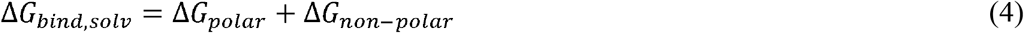

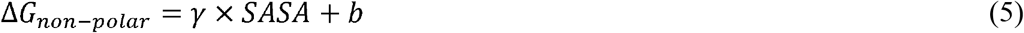

In these equations, Δ*E_MM_*, Δ*G_bind,solv_* and —*T*Δ*S* are calculated in the gas phase. Δ*E_MM_* is the gas phase molecular mechanical energy changes which includes covalent, electrostatic and van der Waals. Based on previous studies the entropy change during binding is small and neglected in these calculations.^44–46^ Δ*G_bind,solv_* is the solvation free energy which comprises the polar and non-polar components. The polar solvation is calculated using the MMPBSA method by setting a value of 80 and 2 for solvent and solute dielectric constants. The non-polar free energy is simply estimated from solvent accessible surface area (SASA) of the solute from equation 5.

## Results

### Structural dynamics

To compute the RMSD of systems, the rotational and translational movements were removed by first fitting the C_α_ atoms of the RBD to the crystal structure and then computing the RMSD with respect to C_α_ atoms of RBD in each system.

Figure 3 shows the RMSD plot in the RBD of SARS-COV, nCOV-2019 and some of its variants. Comparison of RMSD of SARS and nCOV-2019 RBD shows that SARS-COV has a larger RMSD throughout the 500ns simulation. In nCOV-2019, RMSD is very stable with a value of about 1.5 Å whereas in SARS-COV the RMSD increases up to ~4 Å after 100 ns and then fluctuates between 3 and 4 Å. The change in RMSD of SARS is partially related to the motion in the C-terminal which is a flexible loop.

**Figure 3.**
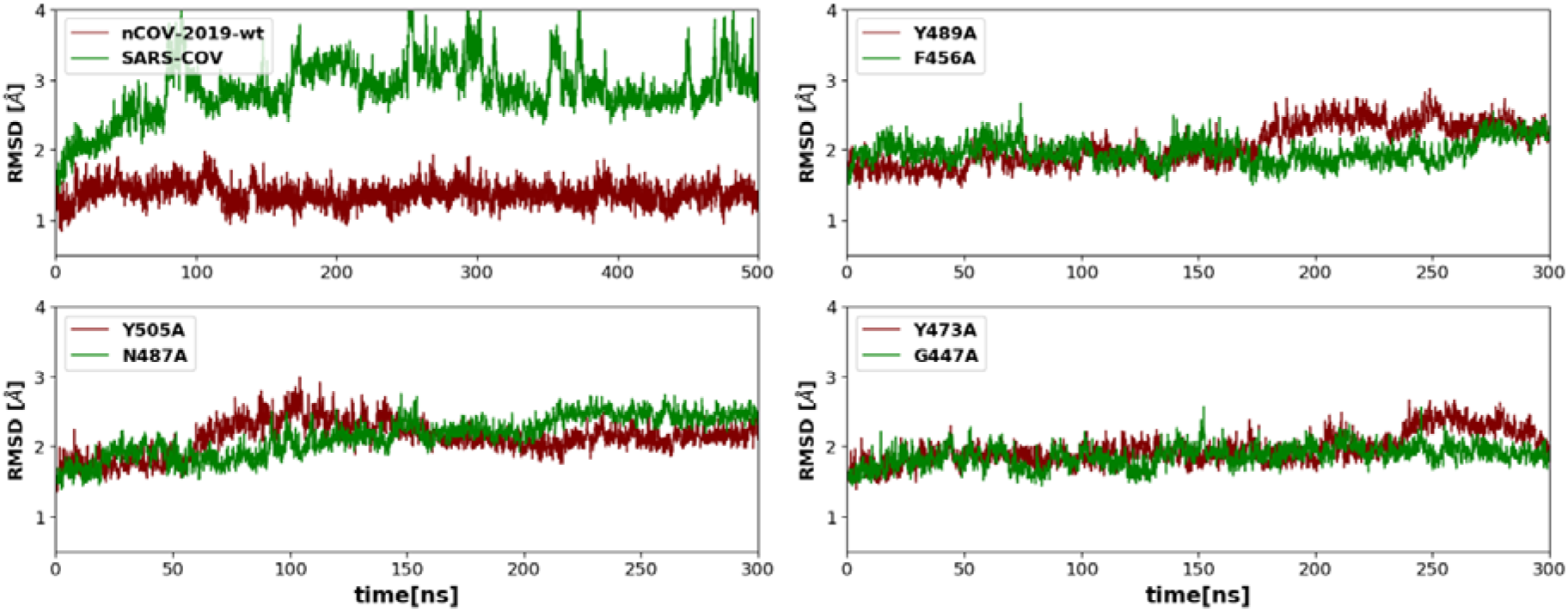
The C_α_ RMSD plots for nCOV-2019 and SARS-COV and a few nCOV-2019 mutations.

The RMSD plots for the nCOV-2019 mutants show similar behaviors to nCOV-2019-wt. In most of the variants, the RMSD is stable during the 300ns simulation. However, a few mutations show some RMSD variance. In mutation Y489A, the RMSD increases from 1.82 ± 0.06 Å to 2.15 ± 0.10 Å after 150 ns. Mutation Y505A resulted in an increase in RMSD up to 100ns to a value to 1.98 ± 0.20 Å and decreased afterwards. RMSD for mutation N487A shows an increasing behavior with a value of 2.10 ± 0.23 Å. In mutation Y473A, the RMSD is 1.94 ± 0.17 Å. Mutations N439K, V483A and V483F showed a stable RMSD of ~1.5 Å. For mutations T478I, G476S, S494P and A475V RMSD increases up to ~2 Å. These variations in the backbone RMSD show the involvement of these residues in the complex stability. RMSD plots for other mutations are shown in Figure S1.

Since the extended loop (residues 449 to 510 shown in figure 1B from *α*_4_ to *α*_5_) of the RBD makes all contacts with ACE2, the RMSD was computed by also fitting to the *C_α_* atoms of this region (Figure S2). The extended loop in nCOV-2019 is very stable with less deviation (RMSD = 0.86 ± 0.017 Å) from the crystal structure compared to SARS-COV having an RMSD of 2.79 ± 0.05 Å. Few of the mutants show an increase in the loop RMSD during the simulation. In mutants N487A and Y449A the loop RMSD jumps to a value of about 2.00 ± 0.04 Å and 1.63 ± 0.09 Å respectively after about 200 ns of simulation. Mutants G447A and E484A show a loop RMSD values of 2.22 ± 0.03 Å and 1.96 ± 0.02 Å during the last 100ns of simulation. The loop RMSD in mutations Y505A, Y473A and S494A is about 1.5 Å during simulation which is higher than nCOV-2019. Other mutants showed a stable extended loop (observed in loop RMSD) during simulation. The loop RMSD for some residues are high which indicates these residues are involved in the stability of the extended loop region.

To characterize of the dynamic behavior for each amino acid in the RBD, we analyzed the root mean square fluctuation (RMSF) of all systems. The RMSF plots for nCOV-2019, SARS-COV and four other mutations are shown in Figure 4. nCOV-2019 shows less fluctuations compared to SARS-COV. *L*_3_ in nCOV-2019 corresponding to residues 476 to 487 (shown in red in Figure 4) has smaller RMSF (1.5 Å) than SARS-COV *L*_3_ residues 463 to 474. *L*_1_ in both nCOV-2019 and SARS-COV (green) has small fluctuation (less than 1.5 Å). Moreover, the C-terminal residues of SARS-COV shows high fluctuation (Figure 4 shown in orange). Few mutants show higher fluctuation in the *L*_1_. Mutants Y505A and S494A had a RMSF of 2.5 Å and mutation N487A has a RMSF of about 4 Å in the *L*_1_ Mutation Y449A has a higher fluctuation RMSF of about 3 Å in the *L*_3_. Mutants G496A, E484A and G447A show a high fluctuation of about 4.5 Å in the *L*_3_. RMSF of other variants are shown in Figure S3.

**Figure 4).**
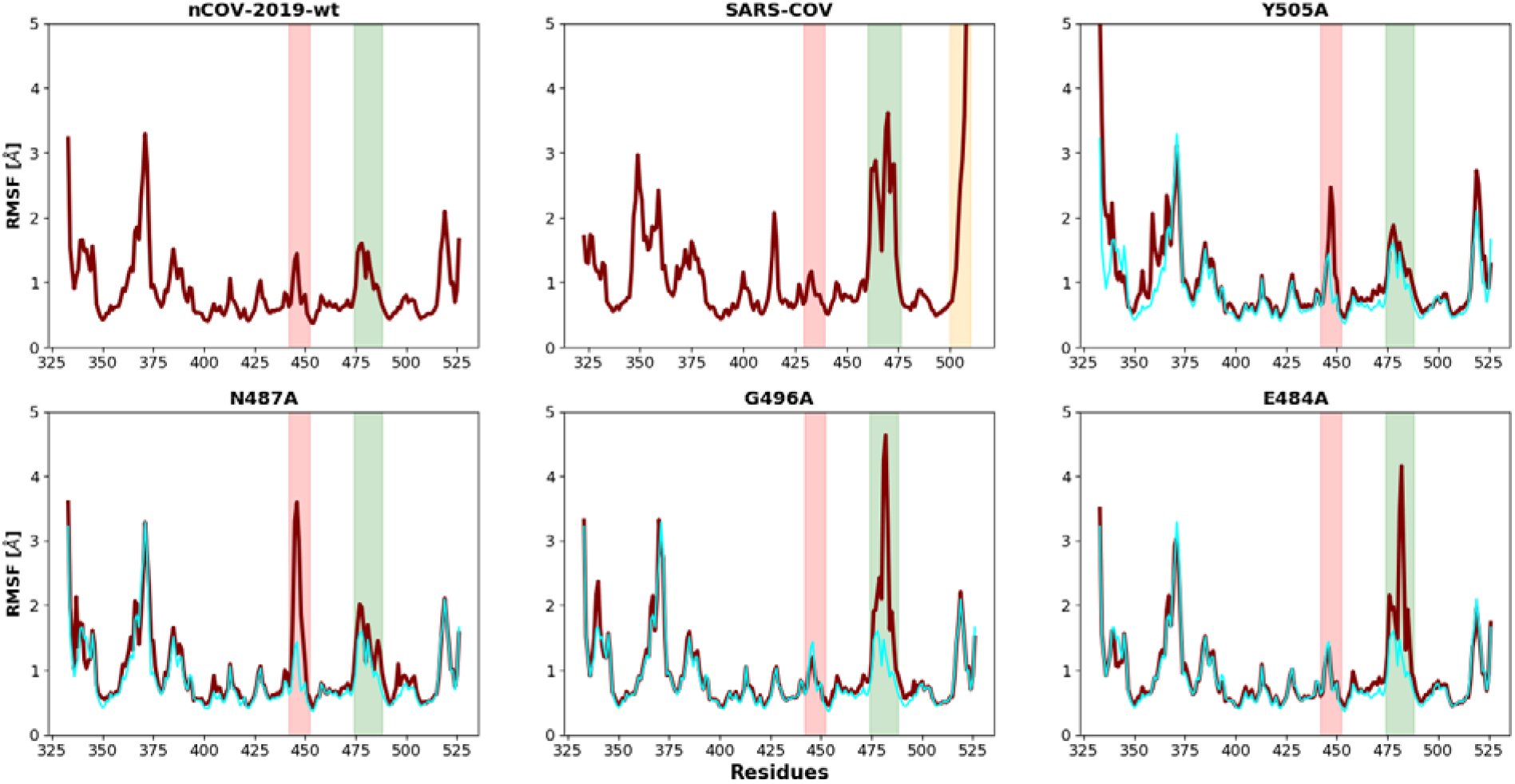
RMSF plots for nCOV-2019-wt, SARS-COV, Y505A, N487A, G496A and E484A. the red shaded region shows the fluctuation in *L*_1_ and the green shaded shows the fluctuation in *L*_3_. The orange shaded region in SARS-COV shows the fluctuation in C-terminal. For comparison, RMSF of nCOV-2019-wt is shown as cyan in other plots.

### Principal component analysis (PCA) and approximate free energy landscape

To identify the dominant motions in the nCOV-2019, SARS-COV, and all the mutants, PCA was performed. Most of the combined motions were captured by the first ten eigenvectors generated from the last 400ns for SARS-COV and nCOV-2019 and the last 200ns for each nCOV-2019 mutant. The percentage of the motions captured by the first three eigenvectors was 51% for nCOV-2019 and 68% for SARS-COV. In all mutations, more than 50% of the motions were captured by the first three eigenvectors. The first few PC’s describe the highest motions in a protein which are related to a functional motion such as binding or unbinding of protein from receptor. The first three eigenvectors were used to calculate the approximate FEL (aFEL) using the last 400ns of simulation for nCOV-2019 and SARS-COV shown in Figure 5, which displays the variance in conformational motion. SARS-COV showed two distinct low free energy states shown as blue separated by a metastable state. There is a clear separation between the two regions by a free energy barrier of about 6-7.5 kcal/mol. These two states correspond to the loop motions in the L_3_ as well as the motion in C-terminal residues of SARS-COV. The *L*_3_ motion in nCOV-2019 is stabilized by H-bond between N487 on RBD and Y83 on ACE2 as well as a *π*-stacking interaction between F486 and Y83. It is evident that the nCOV-2019 is more stable than SASR-COV and exist in one conformation whereas the SARS-COV interface fluctuates and the aFEL is separated into two different regions.

**Figure 5.**
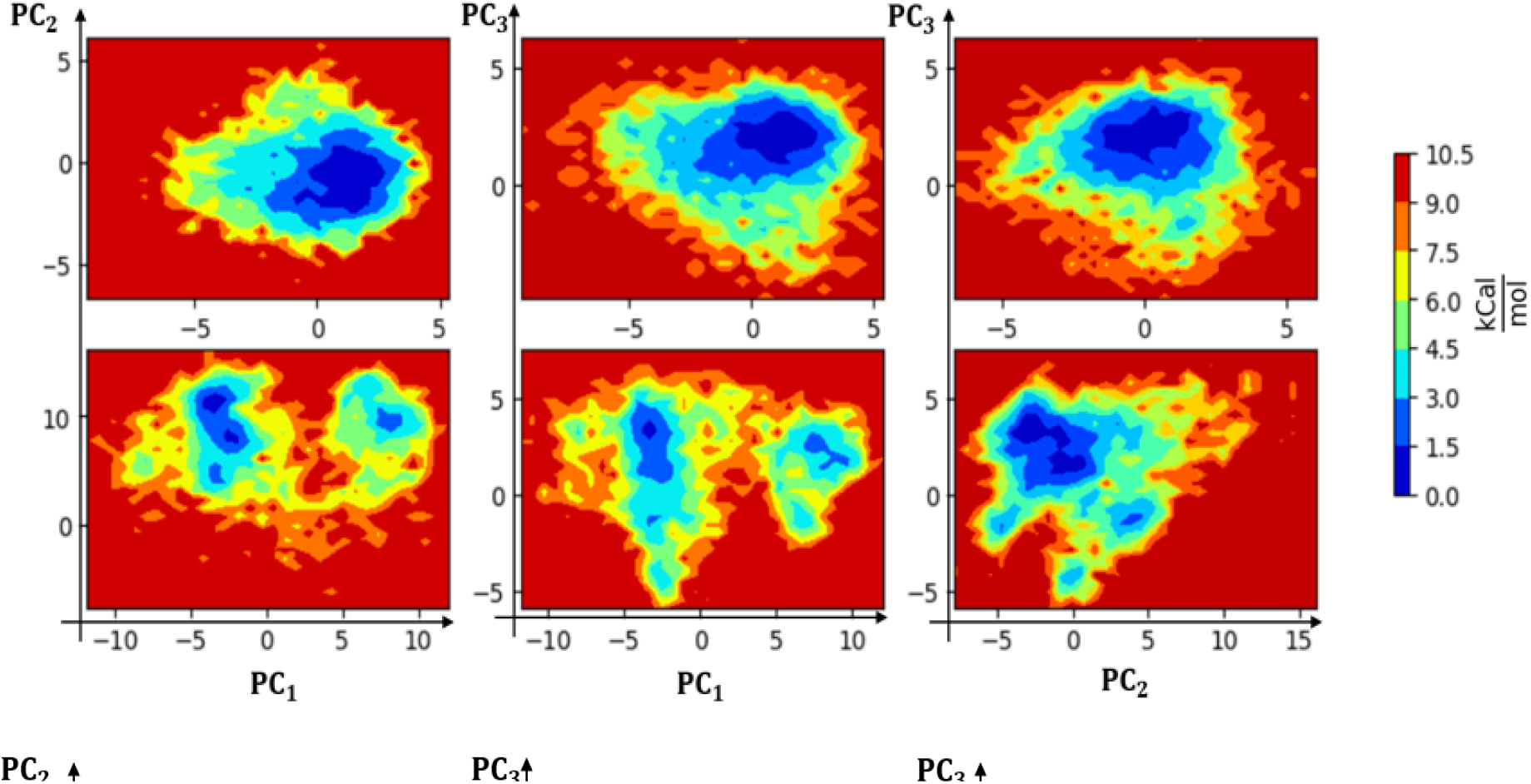
Mapping of the principal components of the RBD for the aFEL from the last 400ns of simulations for SARS-COV (top row) and nCOV-2019-wt (Bottom row). The color bar is relative to the lowest free energy state.

The first two eigenvectors were used to calculate and plot the aFEL as a function of first two principal components using the last 200ns of the simulation for mutant systems. aFEL for a few mutants are shown in Figure 6 and the rest of them are shown in Figure S4. In some mutants such as Q498A and Y453A the aFEL shows only one low energy state. However, aFEL for other mutants such as E484A, G447A, Y449A and N487A separate into multiple regions divided by metastable states. Mutations Y449A, G477A and E484A also showed high fluctutation in *L*_3_ and multiple states in aFEL can be assigned to different states of the *L*_3_. In mutation N487A the *L*_1_ showed large fluctuations and the two states in the aFEL of this mutant correspond to different states of *L*_1_.

**Figure 6.**
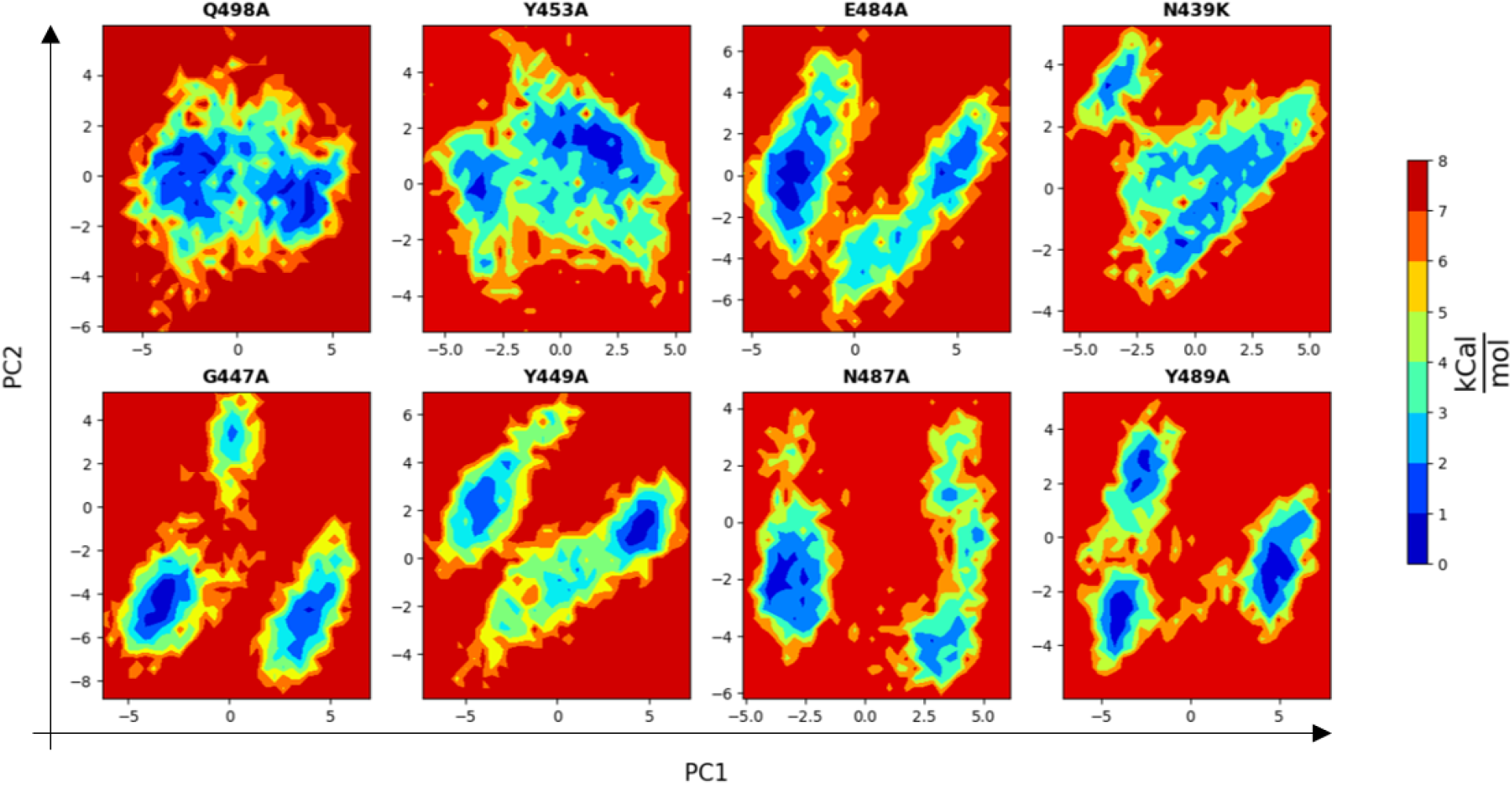
aFEL plots for nCOV-2019, SARS-COV and mutants by projecting onto the first two eigenvectors.

In each system, the first eigenvector was used to construct the porcupine plots to visualize the most dominant movements (Figure 7). nCOV-2019 showed a small motion in L_3_ and the core region and the extended loop region is very rigid showing small cones in the porcupine plot. The core structure of the RBD remains dormant as the cones are blue in most of the regions. In SARS-COV, the C-terminal region shows large motions (Figure 7B). Mutation N487A showed a large motion in *L*_1_ (Figure 7C). Mutations Y449A, G447A and E484A demonstrate large motions in *L*_3_ (Figure 7). Overall, these plots show the involvement of these residues in dynamic stability of RBD/ACE2 complex.

**Figure 7.**
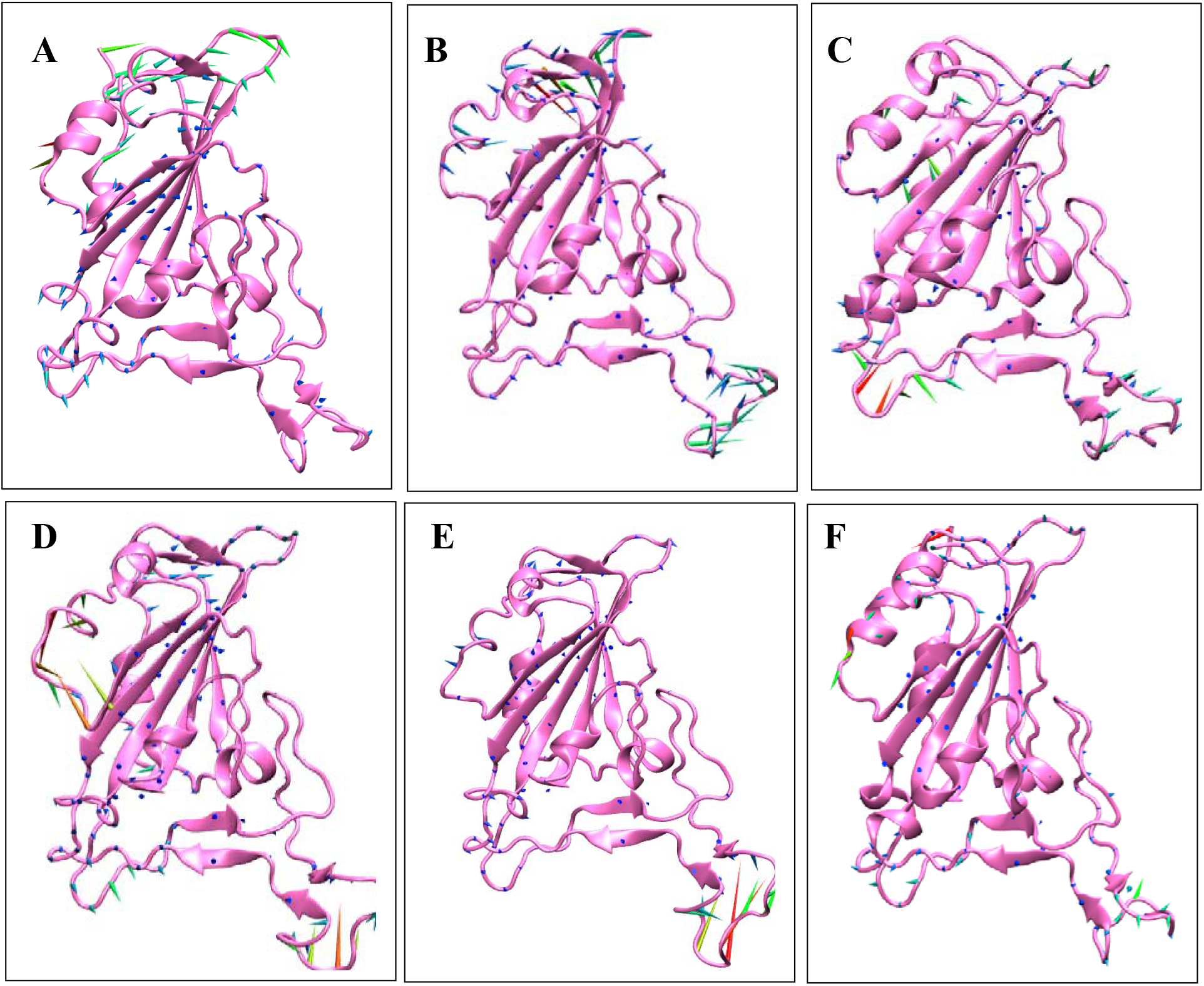
Porcupine plots for A) nCOV-2019 B) SARS-COV C) N487A D) Y449A E) G447A F) 18-Glu484. Arrows show the direction of movement for the *C_α_* atoms and length of arrows show the magnitude of the motion. longer arrows are red and shorter arrows are blue.

### Dynamic cross-correlation maps (DCCM)

The correlated motions of RBD atoms were also analyzed with the DCCM based on the C_α_ atoms of RBD from the last 400ns of simulation for nCOV-2019 and SARS-COV and the last 200ns for the mutant systems (Figure 8). DCCM for nCOV-2019 showed a correlation between residues 490-505 (containing *α*_5_, *L*_4_ and *β*_5_ regions) and residues 440-455 (containing *β*_4_,*L*_1_ and *β*_5_ regions) shown in red rectangle in Figure 8, which is referred to here as *correlation I.* This correlation showed the coordination of these regions for binding ACE2 effectively. Another important correlation that appears in DCCM of nCOV-2019 is between residues 473-481 with residues 482-491. These residues are in *L*_3_ and *β*_6_ regions and their correlation in nCOV-2019 is stronger than SARS-COV. This is due to the presence of *β*_6_ strand in nCOV-2019, whereas in SARS-COV these residues all belong to *L*_3_. This indicates that *L*_3_ in nCOV-2019 has evolved from SARS-COV to adopt new a secondary structure, which causes strong correlation and make the loop act as a recognition region for binding. The correlation in *L*_3_ is shown as blue rectangle in Figure 8. Some of the mutations disrupted the patterns of correlation and anti-correlation in nCOV-2019-wt. Mutation N487A showed a stronger correlation in *L*_3_ and *β*_6_ strand than the wild-type RBD. In mutation E484A, *correlation I* is stronger than the nCOV-2019-wt. DCCM for other mutants are shown in Figure S5. It is worth mentioning that mutation F486A disrupts the DCCM of nCOV-2019 by introducing strong correlations in the core region of RBD as well as the extended loop region. Residue F486A resides in *L*_3_ and plays a crucial role in stabilizing the recognition loop by making a π-stacking interaction with residue Y83A on ACE2.

**Figure 8.**
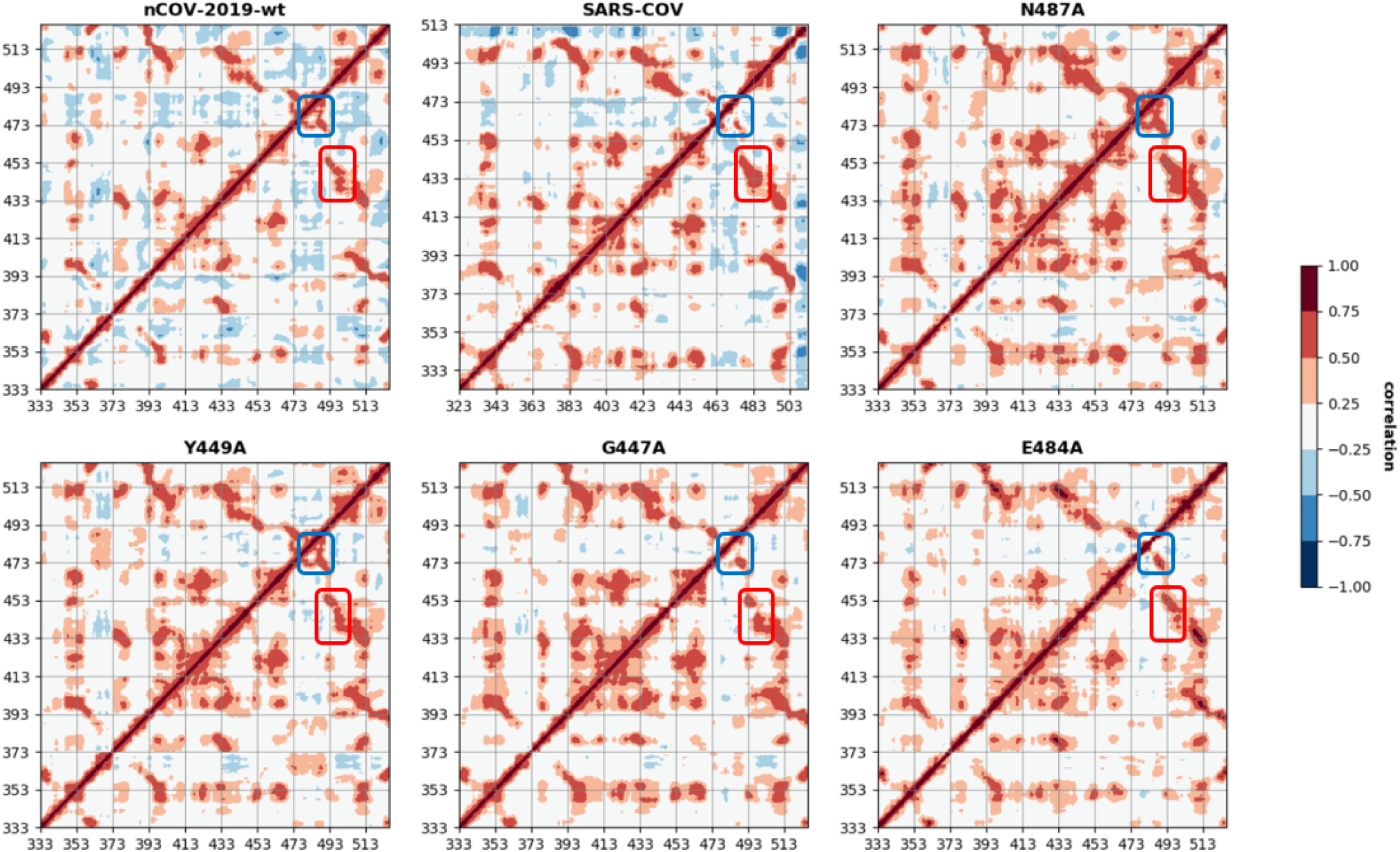
Dynamic cross correlation maps for nCOV-2019, SARS-COV and mutants with residue numbers of the RBD domain. Red box shows the correlation between *α*_5_, *L*_4_ and *β*_5_ regions and *α*_4_, *L*_1_ and *β*_5_. Blue box shows the correlation in *L*_3_ and *β*_6_ regions.

### Binding free energies

The binding energetics between ACE2 and the RBD of SARS-COV, nCOV-2019 and all its mutant complexes were investigated by the MMPBSA method.^44^ The binding energy was partitioned into its individual components including: electrostatic, van der Waals, polar solvation and solvent accessible surface area (SASA) to identify important factors affecting the interface of RBD and ACE2 in all complexes. nCOV-2019 has a total binding energy of 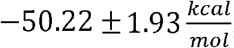, whereas SARS-COV has a much lower binding affinity 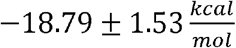. Decomposition of binding energy to its components show that the most striking difference between nCOV-2019 and SARS-COV is the electrostatic contribution which is 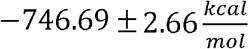 for nCOV-2019 and 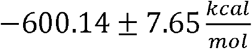 for SARS-COV. This high electrostatic contribution is compensated by a large polar solvation free energy which is 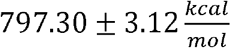 for nCOV-2019 and 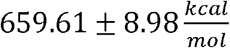 for SARS-COV. nCOV-2019 also possess a higher van der Waals (vdw) contribution 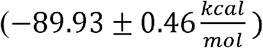 than SARS-COV 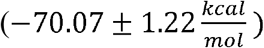. Furthermore, the SASA contribution to binding for SARS-COV was 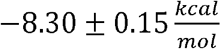 and 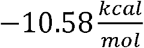 for nCOV-2019.

The binding free energy for nCOV-2019 and SARS-COV was decomposed into a per-residue based binding affinity to find the residues that contribute strongly to the binding and complex formation (Figure 9). Most of the investigated residues in the RBM of nCOV-2019 had a favorable contribution to total binding energy. Residues Q498, Y505, N501, Q493 and K417 contributed more than 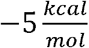 and are crucial for binding. A few residues such as E484 and S494 contributed positively (unfavorable) to the total binding energy. Among all the interface residues K417 had the highest contribution to the total binding energy with 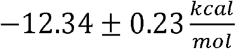. The corresponding residue in SARS-COV is Val404 only had a 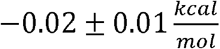 contribution, which points to the importance of this residue for nCOV-2019 binding to ACE2. Residue Q498 contributed to total binding by 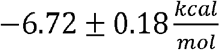 and its corresponding residue in SARS-COV is a Y484 that contributed to total binding by 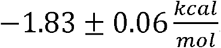. Other important residues Y505 and N501 have more negative contribution to total binding energy than their counterparts in SARS-COV residues Y491 and T487 respectively. Residue D480 in SARS-COV contributed positively to binding energy by 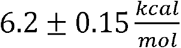 and the corresponding residue in nCOV-2019 which is a S494 residue lowered this positive contribution to only 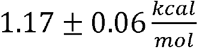. In SARS-COV residue R426 had a high contribution to total binding energy 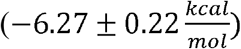 although the corresponding residue in nCOV-2019 is N439 with a contribution of 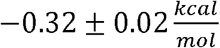. Mutation N439K which is the most observed among all the mutations of RBD can recover part of interaction at this position. These important mutations on RBM of nCOV-2019 from SARS-COV caused RBD of nCOV-2019 to bind ACE2 with much stronger 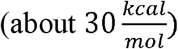 affinity.

**Figure 9.**
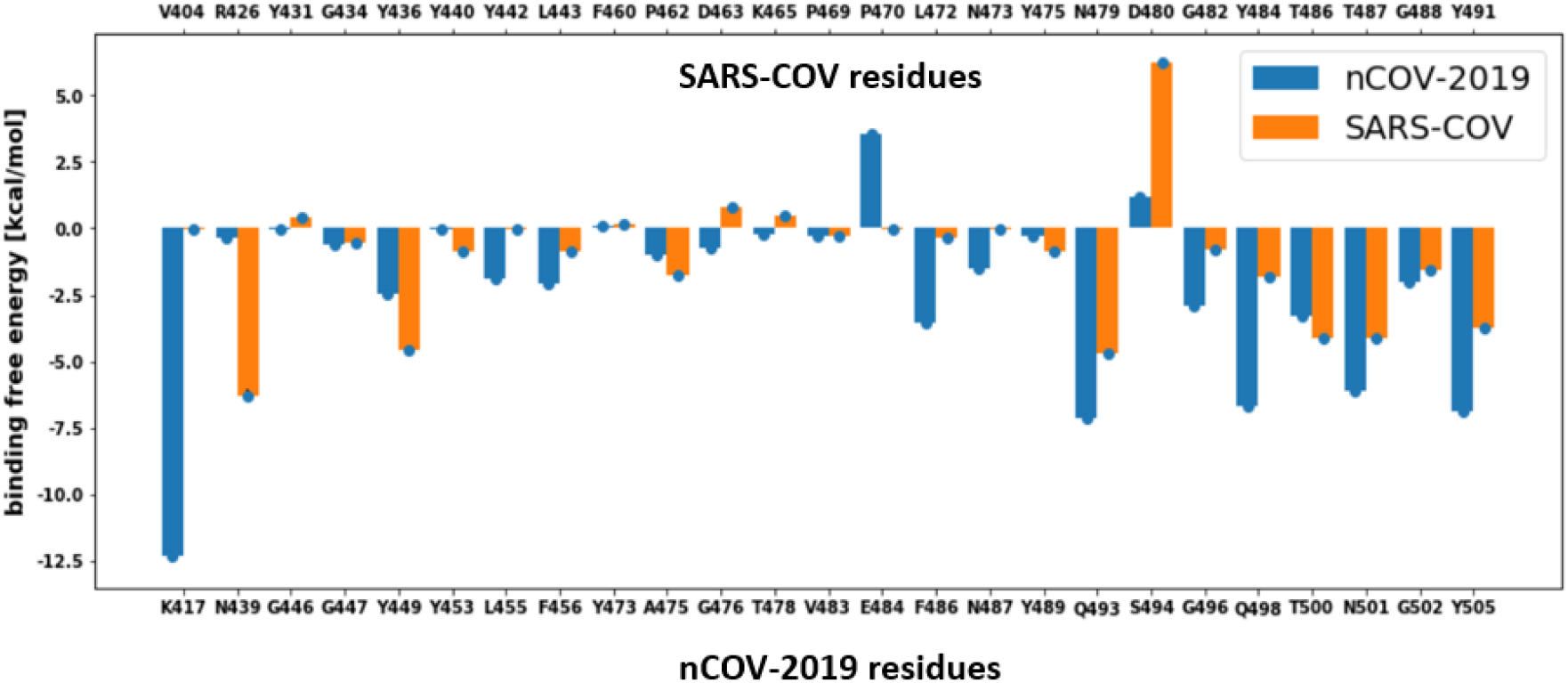
Binding energy decomposition per residue for the RBM of nCOV-2019 and SARS-COV.

Binding free energy decomposition to its individual components for all mutants is represented in Table S2. In all complexes, a large positive polar solvation free energy disfavors the binding and complex formation, which is compensated by a large negative electrostatic free energy of binding. All variants had similar solvent accessible surface area energies which is between —10 and 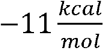 for all complexes. The vdw free energy of binding ranged from 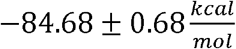 for mutant Q493A to 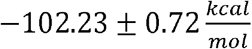 for N487A. Mutant K417A had the lowest electrostatic contribution to binding 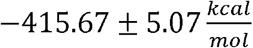 and mutants N439K and E484A had the highest electrostatic binding contribution of 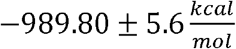 and 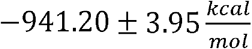 respectively.

Most alanine substitutions exhibited similar or lower total binding energy to nCOV-2019, however a few mutants had higher binding affinity than the wild type. Mutant Y489A had a total binding energy of 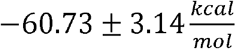 which was about 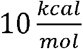 higher than wild type binding energy. Mutants G446A, G447A, Y449A and T478I also demonstrated higher total binding energy than nCOV-2019 with respectively—57.79 ± 2.92, —58.37 ± 2.32, —55.38 ± 2.58 and 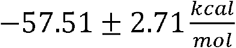 total binding energies. Other alanine substitutions had similar or lower total binding energy than nCOV-2019. Mutant G502A has the lowest binding affinity among all the mutants with a binding energy of 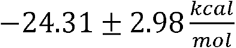. Mutant systems K417A, L455A, T500A and N501A are the other mutant with total binding energies 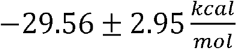 and 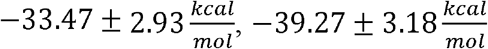 and 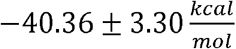 respectively which are significantly lower (less negative) than the wild type complex. The electrostatic component of binding energy contributes the most to this low binding energies for these mutants. The contribution of RBM residues to binding with ACE2 for nCOV-2019 were mapped to the RBD structure and is shown in Figure 11B.

**Figure 10.**
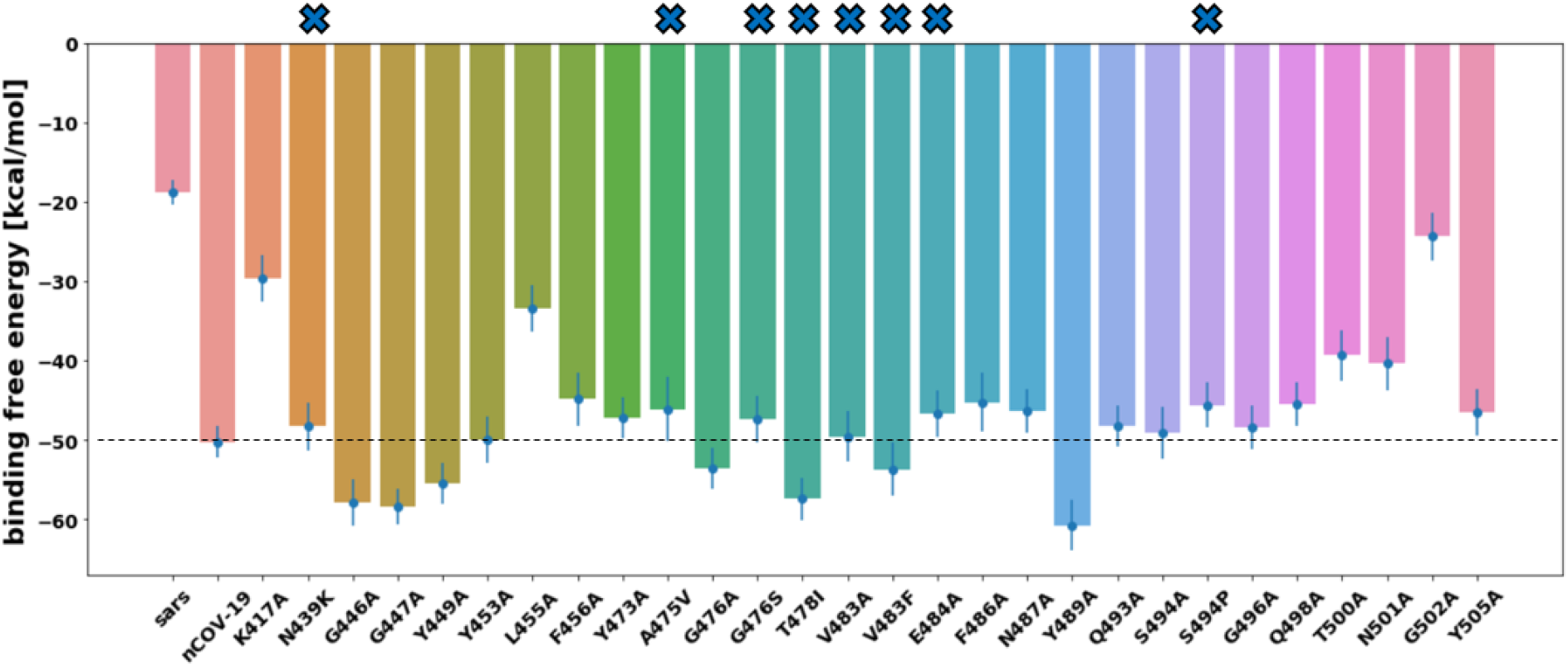
Total free binding energy of SARS-COV, nCOV-2019 and mutants. Natural mutants are shown with X at the bar base.

**Figure 11).**
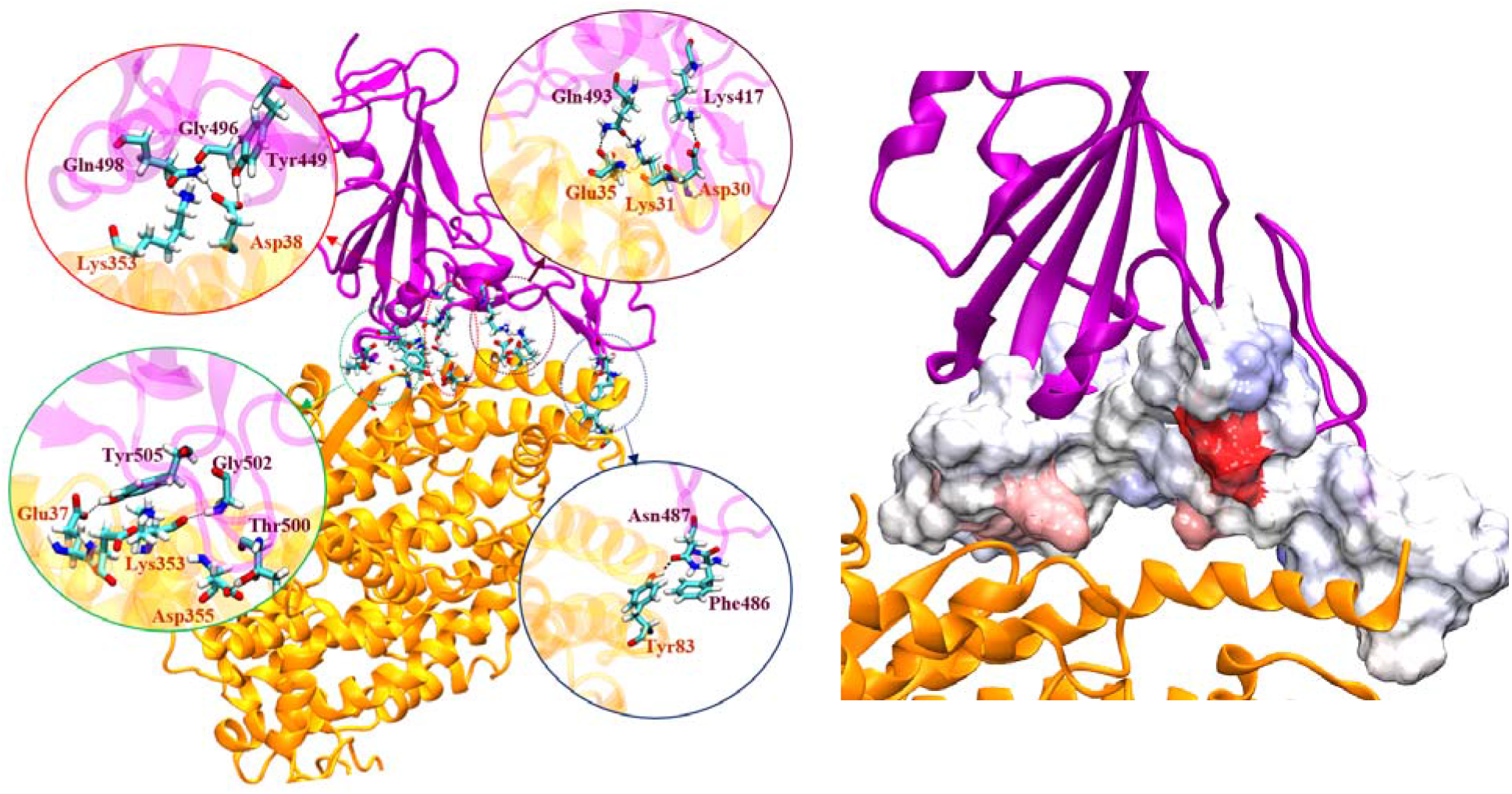
**A)** H bonds between RBD of A) nCOV-2019 and B) SARS-COV. B) mapping contribution of interface residues to structure in RBD of nCOV-2019. The RBD domain is purple and the ACE2 is yellow. The RBD in contact with AC2 is rendered in a surface format with more red being a favorable contribution to binding (more negative) and blue unfavorable contribution (positive).

Natural mutants exhibited similar or higher binding affinities compared to wild-type nCOV-2019. Importantly, mutation T478I which is one of the most occurred mutations in England based on GISAID database^25^ increases the binding affinity to 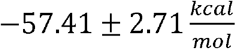 which is about 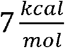 higher than wild-type binding energy. N439K demonstrated a high electrostatic energy which is compensated by large polar solvation energy and this mutant has a total binding energy of 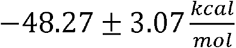. S494P showed a total binding energy of 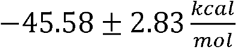 which was lower than the wild type binding energy. Other natural mutants showed binding energies similar to wild-type.

### Hydrogen bond, Salt-bridge and Hydrophobic contact analysis

Important hydrogen bonds (H-bonds) and salt bridges between nCOV-2019 RBD or SARS-COV RBD and ACE2 for the last 400ns of trajectory are shown in Table 1. nCOV-2019 RBD makes 10 H-bonds/1 salt bridge with ACE2 whereas SARS-COV makes only 5 H-bonds/1 salt bridge with ACE2 with more than 30% persistence.

**Table 1.** H-bonds and salt-bridges between nCOV-2019 and ACE2 and SARS-COV and ACE2 that persist for >30%. Salt bridge is shown as bold.

| # | nCOV-2019 | ACE2 | % occupancy | SARS-COV | ACE2 | % occupancy |
| --- | --- | --- | --- | --- | --- | --- |
| 1 | G502 | K353 | 89 | Y436 | D38 | 96 |
| 2 | Q493 | E35 | 83 | **R426** | **E329** | **87** |
| 3 | N487 | Y83 | 80 | T486 | D355 | 83 |
| 4 | Q498 | D38 | 73 | G488 | K353 | 80 |
| **5** | **K417** | **D30** | **55** | N479 | K31 | 52 |
| 6 | T500 | D355 | 53 | Y440 | H34 | 47 |
| 7 | Y505 | E37 | 52 |  |  |  |
| 8 | Q498 | K353 | 49 |  |  |  |
| 9 | Y449 | D38 | 45 |  |  |  |
| 10 | G496 | K353 | 37 |  |  |  |
| 11 | Q493 | K31 | 32 |  |  |  |

The evolution of the coronavirus from SARS-COV to nCOV-2019 has reshaped the interfacial hydrogen bonds with ACE2. G502 in nCOV-2019 has the most persistent H-bond with residue K353 on ACE2. This residue was G488 in SARS-COV, which also makes H-bond with K353 on ACE2. Q493 in nCOV-2019 makes H-bond with E35 and a less persistent H-bond with K31 on ACE2. This residue was an N479 in SARS-COV, which only makes one H-bond with K31 on ACE2 with 52% occupancy. An important mutation from SARS-COV to nCOV-2019 is residue Q498, which was Y484 in SARS-COV. Q498 makes two H-bonds with residues D38 and K353 on ACE2, whereas Y484 in SARS-COV does not make any H-bonds. Importantly a salt bridge between K417 and D30 in nCOV-2019/ACE2 complex contributes to total binding energy by 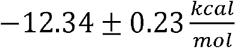. This residue is V404 in SARS-COV which is not able to make any salt-bridge and does not make H-bond with ACE2. Wang et al.^47^ used a free energy perturbation (FEP) approach and showed that mutation V404 to K417 increases the binding affinity of nCOV-2019 RBD to ACE2 by 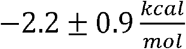. A salt bridge between R426 on RBD and E329 on ACE2 stabilizes the complex in SARS-COV/ACE2. This residue is N439 in nCOV-2019 and is unable to make salt-bridge with ACE2 residue E329. One of the most observed mutations in nCOV-2019 according to GISAID database^25^ is N439K which recovers some of the electrostatic interactions with ACE2 at this position.

The evolution from SARS-COV to nCOV-2019 also has destabilized some specific ACE2/RBD interactions. An important H-bond in SARS-COV is between Y436 on RBD and D38 on ACE2. This residue is Y449 in nCOV-2019 and makes H-bond with D38 on ACE2 but with a lower occupancy than SARS-COV. The unchanged T486 in SARS-COV corresponds to T500 in nCOV-2019, both of which make consistent H-bonds with ACE2 residue D355.

Hydrophobic interactions also play an important role in stabilizing RBD/ACE2 complex in nCOV-2019. An important interaction between nCOV-2019 RBD and ACE2 is the π-stacking interaction between F486 (RBD) and Y83 (ACE2). This interaction helps in stabilizing *L*_3_ in nCOV-2019 compared to SARS-COV where this residue is L472. It was observed by Wang et al. that mutation Leu472 to Phe486 in nCOV-2019 results in a net change in binding free energy of 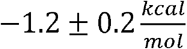. Other interfacial residues in nCOV-2019 RBD that participate in hydrophobic interaction with ACE2 are L455, F456, Y473, A475 and Y489. It is interesting to note that all these residues except Y489 have mutated from SARS-COV. Spinello and coworkers^48^ performed long-timescale (1*μs*) simulation of nCOV-2019/ACE2 and SARS-COV ACE2 and found that *L*_3_ in nCOV-2019 is more stable due to presence of the *β*_6_ strand and existence of two H-bonds in *L*_3_ (H-bonds G485-C488 and Q474-G476). Importantly, an amino acid insertion in *L*_3_ makes this loop longer than *L*_3_ in SARS-COV and enable it to act like a recognition loop and make more persistent H-bonds with ACE2. L455 in nCOV-2019 RBD is important for hydrophobic interaction with ACE2 and mutation L455A lowers the vdw contribution of binding to 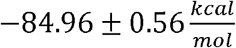 which is about 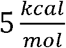 smaller than the wild-type complex. The H-bonds between RBD of nCOV-2019 and SARS-COV are shown in Figure 11A. The structural details discussed here are in agreement with other structural studies of nCOV-2019 RBD/ACE2 complex.^4,49,50^

H-bond analysis was also performed for the mutant systems and the results for H-bonds with more than 40% consistency are in Table S3. Few of the alanine substitutions increase the number of interfacial H-bonds between nCOV-2019 RBD and ACE2. Interestingly, the ala-substitution at Y489A increased the number of H-bonds in the wild-type complex. Mutation in some of the residues having consistent H-bonds in the wild type complex such as Q498A, Q493A and T500A stunningly increased the persistence of H-bonds in the neighboring residues and maintaining the number of H-bonds in the wild-type complex. This indicates that the RBM of nCOV-2019 can reshape the network of H-bonds and strengthen the H-bonds upon mutation in these locations. However, few mutations also decrease the number of H-bonds from the wildtype complex. Alanine substitution at residue G502 has a significant effect on the network of H-bonds between nCOV-2019 and SARS-COV. This residue locates at the end of *L*_4_ loop near two other important residues Q498 and T500. This mutation breaks the H-bonds at these residues. Mutation K417 decreases the number of H-bonds to only 5 where the H-bond at residue Q498 is broken. This indicates the delicacy of H-bond from residue Q498 which can easily be broken upon ala-substitution at other residues. Furthermore, mutation N487 also decreases the number of H-bonds by breaking the H-bond at Q498.

## Discussion

In this work, we preformed MD simulations to unveil the detailed molecular mechanism for receptor binding of nCOV-2019 and comparing it with SARS-COV. The role of key residues at the interface of nCOV-2019 with ACE2 were investigated by computational ala-scanning. A rigorous 500ns MD simulation was performed for nCOV-2019 and SARS-COV as well as 300ns MD simulation on each mutant. These simulations aiding in our understanding in the dynamic role of RBD/ACE2 interface residues and estimating the binding free energy of these variants, which shed light on crucial residues for the RBD/ACE2 complex stability. Moreover, numerous mutations have been identified in the RBD of different nCOV-2019 strains from all over the world not known to be critical for infection.^51^ The role of these mutations on the stability of RBD/ACE2 complex were investigated to shed light on their role in the viral infection of coronavirus.

Changes in RBD structure of nCOV-2019, SARS-COV and mutants from their crystal structure were analyzed by RMSD and RMSF. nCOV-2019 showed a stable structure with a RMSDU) Å, whereas SARS-COV had a larger RMSD value between 3-4 Å during the simulation. Most mutations of nCOV-2019 maintained similar stability to the wild-type. However, in few nCOV-2019 mutations resulted in larger deviations (>2Å), i.e., Y489A, F456A, Y505A, N487A, K417A, Y473A, Y449A.. We further investigated the structure of the extended loop domain (Figure 1B) and discovered that nCOV-2019 is stable with RMSD of less than 1 *Å*, whereas the extended loop in SARS-COV shows an RMSD of about 3Å during simulation (figure S2). Some mutants showed high RMSD in this region. Alanine-substitution at residue N487 increased the extended loop RMSD to 2.5 Å. Other mutations that increased the extended loop RMSD (>2 Å) include Y449A, G477A and E484A. The dynamic behavior of RBD was further investigated by analyzing the RMSF of all systems. As shown in Figure 4, nCOV-2019 shows less fluctuation in *L*_3_ than SARS-COV. This is due to the presence of a 4-residue motif (GQTQ) in nCOV-2019 *L*_3_, which forces the loop to adopt a compact structure by making 2 H-bonds (G485-C488 and Q474-G476) and thereby reducing the fluctuations in the loop. Residues F486 and N487 play major roles in stabilizing the recognition loop by making π-stacking and H-bond interactions with residue Y83 on ACE2. Alanine substitution at N487 introduced large RMSF to *L*_1_. Mutation L472 to F486 was shown to favor binding by 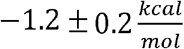 using FEP^47^. In addition, this mutation was shown to be among the five mutations that produce a super affinity ACE2 binder based on SARS-COV RBD.^6^ Alanine mutations at residues Y449, G447 and E484 increased the motion in *L*_3_ characterized by large RMSF in this region.

Using principal component analysis, the free energy landscape for nCOV-2019 and SARS-COV demonstrated that the former occupies only one low energy state whereas the latter forms two distinct low energy basins separated by a metastable state with a barrier of about 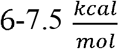. This confirms that the level of binding for the RBD domain is weaker in SARS-COV due to the presence of two basins. Similarly, alanine-substitution for a few residues (Y489A, N487A, Y449A and E484A) caused the free energy landscape to degenerate into separate multiple low energy regions. Dominant motions in the RBD are visualized in Figure 7 using the first eigenvector of the PC A. nCOV-2019 and SARS-COV did not show any strong motion in the extended loop region. Porcupine plots of alanine-mutants demonstrated that mutant N487A shows large motion in *L*_1_ region and Y449A, G447A and E484A showed large motions in *L*_3_.

To better characterize the functional motions of RBD, DCCM for all systems are constructed and demonstrated in Figure 8. nCOV-2019 showed large correlation between the *α*_4_-*L*_1_-*β*_5_ and *α*_5_-*L*_4_-*β*_5_ region. This correlation was stronger in SARS-COV and few mutants such as Y449A, G447A and E484A. Another important correlation in nCOV-2019 is inside *L*_3_ and *β*_6_. This correlation is stronger in nCOV-2019 than SARS-COV due to the presence of *β*_6_ which makes the loop to adopt correlated motions. Few mutants impact the correlation in this region such as N487A. DCCM maps for other mutants are presented in Figure S5. Interestingly, mutant F486A which resides in *L*_3_ and participates in binding by π-stacking interaction with Y83 on ACE2, disrupts the DCCM of wild-type nCOV-2019 and introduces strong correlation in the extended loop region as well as core structure of RBD.

The details of hydrogen bond and salt-bridge pattern in nCOV-2019 and SARS-COV to ACE2 (Table 1) are key to the virus attachment to the host. nCOV-2019 residues participate in 10 H-bonds/1 salt bridge with ACE2, whereas SARS-COV only has 5 H-bonds/1 salt bridge with ACE2. This significantly contributes to the 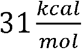 difference in total binding free energy of nCOV-2019 and SARS-COV. The binding energies calculated here for nCOV-2019 and SARS-COV (—50.22 ± 1.93 and 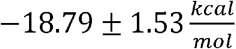, respectively) are in good agreement with the binding energies calculated using Generalized Born method (GB) by Spinello et al.^52^ Moreover, the patterns of H-bonds between nCOV-2019 and ACE2 was also already characterized by other groups^47,52^, which agrees with our work. An important H-bond between nCOV-2019 and ACE2 is between G502 on RBD and K353 of ACE2. G502 is in *L*_4_ region, which is populated by 5 H-bonds between RBD and ACE2. The contribution of this residue to the total binding energy is 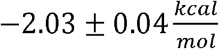 and the Ala-substitution at G502 has the highest effect on the binding energy among all the residues by reducing the total binding energy to 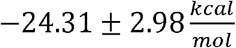 which is the lowest among all mutants. This mutation breaks the other H-bonds in *L*_4_ such as H-bonds from residues Q498 and T500. This residue is preserved and corresponds to residue G488 in SARS-COV, which also makes a persistent H-bond with residue K353 on ACE2. Residue Q493 in nCOV-2019 participates in binding ACE2 by making two H-bonds with residues E35 and K31 on ACE2. Q493 corresponds to residue N479 in SARS-COV, which only makes one H-bond with residue Lys31 on ACE2 with low consistency (52%). This caused Q493 to have more contribution to total binding 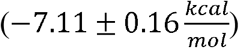 than its counterpart N479 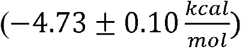. However, alanine substitution at Q493 did not have a large effect on total binding energy and this mutant had a total binding energy of 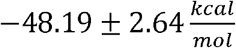. This mutation increases the persistence of H-bonds in neighboring residues and maintains the number of persistent H-bonds in the wild-type complex. Residues Q498 and T500 in nCOV-2019 are crucial for binding by making H-bonds with ACE2 residues D38, D355 and K353. Residue Q498 corresponds to residue Y484 in SARS-COV which does not make any H-bond in SARS-COV/ACE2 complex. Q498 contributes to binding by 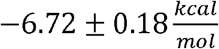 which is more than the contribution of Y484 in SARS-COV 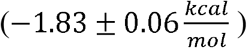. Ala-substitution at Q498 did not show large impact on total binding energy. Residue T500 is conserved and corresponds to residue T486 which also makes a H-bond with Asp355 on ACE2. Mutation of T500 to Alanine lowers the binding energy to 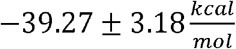, which is about 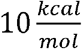 lower than the wild-type complex. Residue N487 in nCOV-2019 locates in *L*_3_ and plays a crucial role in stabilizing the recognition loop by making H-bond with Y83 on ACE2. This residue contributes to total binding energy of nCOV-2019 by 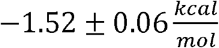, whereas its corresponding residue in SARS-COV does not show any contribution to binding energy 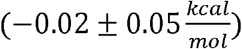. This demonstrates that *L*_3_ in SARS-COV has evolved to be an important recognition loop in nCOV-2019, which participates in binding with ACE2. Residue K417 in nCOV-2019 has the most contribution to total binding energy 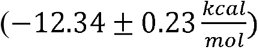 by making a salt-bridge with residue D30 on ACE2. This residue is crucial for binding of RBD and ACE2 and alanine substitution lowers the total binding energy to 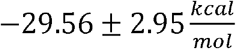. This salt-bridge is found to be important for stability of the crystal structure of RBD/ACE2 complex in nCOV-2019.^50^ This residue is Val404 in SARS-COV which does not participate in binding ACE2. Another important residue in nCOV-2019 is L455 which contributes to binding by 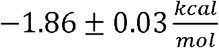. This residue is important for hydrophobic interaction with ACE2 and mutating it to alanine lowers the total binding energy to 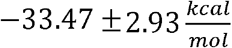. The vdw energy for this mutant is about 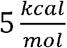 lower than the wild-type complex, which is part of the reason for lowering the total binding energy of this mutant.

Total binding energy calculation of all the variants showed that mutation Y489A has the strongest binding energy among all systems which is about 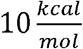 stronger than that of nCOV-2019 complex. This residue is located in *β*_6_, which is part of the recognition loop. This mutation causes the extended loop to move closer to ACE2 interface and all the H-bonds in the wild-type complex are more consistent after alanine substitution at this location. Per-residue binding energy decomposition showed that this residue participates in binding only by 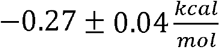. A high electrostatic interaction energy of 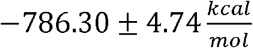 is the reason for higher binding energy of mutant Y489A than wild-type complex. It is interesting to note that among the 5 residues L455, F456, Y473, A475 and Y489 that make hydrophobic interactions with ACE2, Y489 is the only residue that is conserved from SARS-COV. This shows the capacity of the virus to evolve even further in the RBD to bind the receptor ACE2 with stronger affinity. Other mutations that increase the binding energy are G446A, G447A, Y449A and T478I. Residues G446, G447 and Y449 all reside in *L*_1_ and mutation to alanine can make *L*_1_ take a more rigid form as shown in RMSF plot in Figure S3.

Important mutations found in naturally-occurring nCOV-2019 appear to influence to some extent the binding to ACE2. Mutation T478I which is one of the highly seen mutations in England according to GISAID database, increases the binding energy of nCOV-2019 to ACE2 by about 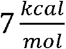. Mutation N439K has the highest occurrence among all strains of coronavirus in the GISAID database which demonstrated the highest electrostatic interaction among all studied systems. This residue corresponds to R426 in SARS-COV which exerts a salt-bridge interaction with E329 on ACE2. Mutation N439K recovers some of this ACE2 interaction. Mutant E484A which is also one of the observed mutations in Spain based on GISAID database, demonstrates a high electrostatic interaction with ACE2. E484 contributes to binding by 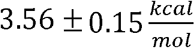 whereas the corresponding residue in SARS-COV, P469 contributes to binding of SARS-COV to ACE2 by 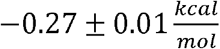. This residue is close to D30 on ACE2 and have electrostatic repulsion with this residue. High electrostatic energy in mutations N439K and E484A are compensated by a large polar solvation energy and these mutants show total binding energies similar to wild-type complex. This could be due to the use of MMPBSA approach for calculation of polar solvation and further studies are needed to study the effect of these mutations on viral infectivity of coronavirus.

Additional sequence differences between nCOV-2019 and SARS-COV influence RBD/ACE2 binding. Residue D480 in SARS-COV contributes negatively to total binding energy 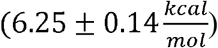 and mutation of this residue to S494 in nCOV-2019 lowers this negative contribution to 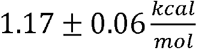. D460 in SARS-COV is located in a region of high negative charge from residues E35, E37 and D38 on ACE2. Electrostatic repulsion between D480 on SARS-COV and the acidic residues on ACE2 is the reason for highly negative contribution of this residue to binding of SARS-COV to ACE2. Mutation to S494 in this location removes this highly negative contribution.

To our knowledge this is first detailed molecular simulation study on the effect of mutations on binding of nCOV-2019 to ACE2. Previous computational studies have found that nCOV-2019 binds to ACE2 with a total binding affinity which was about 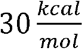 stronger than SARS-COV and is in fair agreement with the results here.^52^ The critical role of interface residues and residues are computationally investigated here and in other articles and the results of all the studies indicate the importance of these residues for the stability of the complex and finding hotspot residues for the interaction with receptor ACE2.^47,52–54^ It is interesting to note the role of *L*_3_ in the stability of the RBD/ACE2 complex. The amino acid insertions in *L*_3_ for nCOV-2019 has converted as unessential part of RBD in SARS to a functional domain of the RBD. This loop participates in binding ACE2 by making H-bond as well as *π*-stacking interaction with ACE2, which makes this region to act as a recognition loop. Previous studies on SARS-COV have shown there is a correlation between higher binding affinity to receptor and higher infection rate by coronavirus.^55–58^ High binding affinity for some mutants such as T478I could be the reason for higher human-to-human transmission rate in regions where these mutations are found. It is also alerting that mutations at other residues such as G446A, G449A, Y449A and Y489A increase the binding affinity considerably and should be monitored. Mutations of nCOV-2019 RBD that do not change the binding affinity and complex stability, could have implications for antibody design purposes since they could act antibody escape mutants. Escape from monoclonal antibodies are observed for mutations of SARS-COV in 2002 and these mutations should be considered for any antibody design endeavors against consider these escape mutations.

## Conclusion

In conclusion, this study unraveled key molecular traits underlying the higher affinity of nCOV-2019 for ACE2 compared to SARS-COV and unveiled critical residues for the interaction by *in-silico* alanine scanning mutations and binding free energy calculations. The higher affinity of nCOV-2019 to binding with ACE2 correlates with higher human-to-human transmissibility of nCOV-2019 compared to SARS-COV. Ala-scanning mutagenesis of the interface residues of nCOV-2019 RBM has shed light on the crucial interface residues and helped obtain an atomic-level understanding of the interaction between coronavirus and the receptor ACE2 on the host cell. MD simulations on RBD mutations found in strains of nCOV-2019 from different countries aiding in the understanding of how these mutations can play and importance to in viral infection with ACE2 attachment. In addition to previously reported residues, it was found that residue F486 locating in *L*_3_ plays a crucial role in dynamic stability of the complex by a *π*-stacking interaction with ACE2. Per-residue free energy decomposition pinpoints the critical role of residues K417, Y505, Q498, Q493 in binding ACE2. Alanine scanning of interface residues in nCOV-2019 RBD showed that alanine substitution at some residues such as G502, K417 and L455 can significantly decrease the binding affinity of the complex. Moreover, mutation T478I, which is one of the most probable mutations in RBD of nCOV-2019 is found to binding ACE2 with about 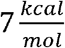 higher affinity than wild-type. Other occurring mutations N439K and E484A are found to increase the electrostatic interaction of RBD with ACE2. It is also alerting that some of the alanine substitutions at residues G446, G447 and Y489 substantially increased the binding affinity that may lead to a strongly RBD attachment to ACE2 and influence the infection virulence. On the other hand, most mutations are found not to impact the binding affinity of RBD with ACE2 in nCOV-2019 which could have implications for vaccine design endeavors as these mutations could act as antibody escape mutants. Receptor recognition is the first line of attack for coronavirus and this study gives novel insights to key structural features of interface residues for advancement of effective therapeutic strategies to stop the coronavirus pandemic.

## Dedication

The authors would like to dedicate this article to the doctors and nurses who sacrificed their time, health and even their lives to fight COVID-19, particularly those in Iran and the United States. JBK would also like to dedicate this work to family friend Joe Kaplan (Silver Spring, MD) who passed away due to COVID-19 on April 22, 2020.

## Acknowledgement

This work was supported by National Heart, Lung and Blood institute (NHLBI) at the National Institute of Health (NIH). The authors acknowledge the biowulf high performance computation center for providing the time and resources for this project.

