## Supplemental Fig/Tables for "Critical Sequence Hot-spots for Binding of nCOV-2019 to ACE2 as Evaluated by Molecular Simulations"


**Table S1.** Residues investigated here and their corresponding location in RBD of nCOV-2019

| residue | Location |
|---|---|
| **K417** | $\alpha_3$ |
| **N439** | $\alpha_4$ |
| **G446** | $L_1$ |
| **G447** | $L_1$ |
| **Y449** | $L_1$ |
| **Y453** | $\beta_5$ |
| **L455** | $\beta_5$ |
| **F456** | $L_2$ |
| **Y473** | $\beta_6$ |
| **A475** | $\beta_6$ |
| **G476** | $L_3$ |
| **T478** | $L_3$ |
| **V483** | $L_3$ |
| **E484** | $L_3$ |
| **F486** | $L_3$ |
| **N487** | $L_3$ |
| **Y489** | $\beta_6$ |
| **Q493** | $\beta_5$ |
| **S494** | $\beta_5$ |
| **G496** | $L_4$ |
| **Q498** | $L_4$ |
| **T500** | $L_4$ |
| **N501** | $L_4$ |
| **G502** | $L_4$ |
| **Y505** | $\alpha_5$ |

**Table S2.** Binding free energy decomposition in $\frac{kcal}{mol}$ for 2019-nCOV, SARS-COV and mutants

| | vdw | Electro | Polar solv | SASA | total |
|---|---|---|---|---|---|
| **SARS-COV** | -70.07 ± 1.22 | -600.14 ± 7.65 | 659.61 ± 8.98 | -8.39 ± 0.15 | -18.79 ± 1.53 |
| **nCOV-2019** | -89.93 ± 0.46 | -746.59 ± 2.66 | 797.3 ± 3.12 | -10.58 ± 0.05 | -50.22 ± 1.93 |
| **K417A** | -88.23 ± 0.58 | -415.67 ± 5.07 | 484.87 ± 4.89 | -10.29 ± 0.09 | -29.56 ± 2.95 |
| **N439K** | −95.4 ± 0.63 | −989.84 ± 5.57 | 1047.70 ± 5.08 | −10.86 ± 0.06 | −48.27 ± 3.07 |
| **G446A** | -91.9 ± 0.5 | -730.12 ± 3.68 | 774.7 ± 4.18 | -10.6 ± 0.08 | -57.79 ± 2.92 |
| **G447A** | -93.95 ± 0.75 | -756.61 ± 4.63 | 803.48 ± 5.02 | -11.08 ± 0.09 | -58.37± 2.32 |
| **Y449A** | -98.13 ± 0.78 | -721.06 ± 4.69 | 775.51 ± 4.38 | -10.82 ± 0.07 | -55.38 ± 2.58 |
| **Y453A** | -92.3 ± 0.65 | -712.76 ± 3.61 | 765.63 ± 3.98 | -10.96 ± 0.07 | -49.98 ± 2.92 |
| **L455A** | -84.96 ± 0.56 | -734.72 ± 4.44 | 795.41 ± 4.06 | -10.63 ± 0.08 | -33.47 ± 2.93 |
| **F456A** | -95.43 ± 0.57 | -770.12 ± 3.38 | 832.56 ± 4.14 | -11.59 ± 0.07 | -44.84 ± 3.38 |
| **Y473A** | -90.48 ± 0.54 | -725.17 ± 4.27 | 779.23 ± 4.16 | -10.61 ± 0.06 | -47.23 ± 2.59 |
| **A475V** | −93.12 ± 0.49 | −712.12 ± 4.59 | 769.23 ± 4.71 | −10.68 ± 0.08 | −46.07 ± 4.11 |
| **G476A** | -92.84 ± 0.57 | -746.46 ± 4.92 | 796.97 ± 4.84 | -10.99 ± 0.08 | -53.57 ± 2.58 |
| **G476S** | −92.25 ± 0.64 | −712.08 ± 4.10 | 767.40 ± 4.32 | −10.77 ± 0.08 | −47.38 ± 2.86 |
| **T478I** | −89.58 ± 0.84 | −756.13 ± 4.39 | 799.32 ± 3.78 | −10.64 ± 0.08 | −57.41 ± 2.71 |
| **V483A** | −90.74 ± 0.60 | −737.48 ± 3.94 | 789.22 ± 3.74 | −10.76 ± 0.07 | −49.55 ± 3.17 |
| **V483F** | −87.23 ± 0.6 | −738.08 ± 4.73 | 782.82 ± 4.04 | −10.59 ± 0.09 | −53.70 ± 3.35 |
| **E484A** | -95.7 ± 0.62 | -941.2 ± 3.95 | 1002.21 ± 4.92 | -11.15 ± 0.08 | -46.67 ± 2.89 |
| **F486A** | -90.07 ± 0.66 | -724.64 ± 4.39 | 779.73 ± 5.01 | -10.68 ± 0.09 | -45.23 ± 3.66 |
| **N487A** | -102.23 ± 0.72 | -724.44 ± 3.69 | 791.41 ± 4.3 | -11.37 ± 0.08 | -46.33 ± 2.73 |
| **Y489A** | -89.49 ± 0.69 | -786.3 ± 4.74 | 826.66 ± 4.53 | -10.75 ± 0.08 | -60.73 ± 3.14 |
| **Q493A** | -84.68 ± 0.68 | -713.28 ± 3.67 | 758.56 ± 3.63 | -9.87 ± 0.07 | -48.19 ± 2.64 |
| **S494A** | -94.98 ± 0.66 | -736.93 ± 4.05 | 793.94 ± 3.84 | -11.02 ± 0.07 | -49.09 ± 3.26 |
| **S494P** | −86.40 ± 0.53 | −712.25 ± 4.76 | 766.55 ± 3.98 | −10.29 ± 0.08 | −45.58 ± 2.83 |
| **G496A** | -93.17 ± 0.55 | -728.93 ± 4.39 | 784.81 ± 4.93 | -10.89 ± 0.07 | -48.38 ± 2.67 |
| **Q498A** | -90.48 ± 0.61 | -756.18 ± 4.6 | 812.84 ± 4.75 | -11.02 ± 0.09 | -45.4 ± 2.74 |
| **T500A** | -93.64 ± 0.62 | -704.44 ± 4.1 | 769.65 ± 4.73 | -10.86 ± 0.08 | -39.27 ± 3.18 |
| **N501A** | -88.59 ± 0.66 | -730.53 ± 3.63 | 788.41 ± 4.3 | -10.75 ± 0.08 | -40.36 ± 3.3 |
| **G502A** | -87.61 ± 0.63 | -706.13 ± 4.56 | 780.08 ± 4.61 | -10.51 ± 0.07 | -24.31 ± 2.98 |
| **Y505A** | -91.35 ± 0.7 | -746.12 ± 4.31 | 802.16 ± 5.29 | -10.78 ± 0.08 | -46.49 ± 2.92 |

**Table S3.** Details of H-bonds for mutants of nCOV-2019

| mutant | # | RBD | ACE2 | Occupancy % |
|---|---|---|---|---|
| K417A | 1 | GLY502 | LYS353 | 87 |
| | 2 | ASN487 | TYR83 | 82 |
| | 3 | GLN493 | GLU35 | 77 |
| | 4 | THR500 | ASP355 | 73 |
| | 5 | TYR449 | ASP38 | 53 |
| N439K | 1 | GLY502 | LYS353 | 85 |
| | 2 | GLN493 | GLU35 | 82 |
| | 3 | ASN487 | TYR83 | 67 |
| | 4 | GLN498 | ASP38 | 66 |
| | 5 | THR500 | ASP355 | 60 |
| | 6 | LYS417 | ASP30 | 51 |
| | 7 | GLN498 | LYS353 | 42 |
| G446A | 1 | TYR449 | ASP38 | 94 |
| | 2 | GLY502 | LYS353 | 90 |
| | 3 | GLN498 | ASP38 | 88 |
| | 4 | TYR83 | ASN487 | 79 |
| | 5 | GLN493 | GLU35 | 76 |
| | 6 | THR500 | ASP355 | 54 |
| | 7 | LYS353 | GLN498 | 51 |
| | 8 | LYS353 | GLY496 | 46 |
| | 9 | TYR505 | GLU37 | 43 |
| G447A | 1 | TYR449 | ASP38 | 92 |
| | 2 | GLY502 | LYS353 | 81 |
| | 3 | GLN493 | GLU35 | 68 |
| | 4 | TYR505 | GLU37 | 57 |
| | 5 | TYR83 | ASN487 | 53 |
| | 6 | THR500 | ASP355 | 46 |
| | 7 | LYS353 | GLN498 | 42 |
| Y449A | 1 | GLY502 | LYS353 | 84 |
| | 2 | GLN493 | GLU35 | 78 |
| | 3 | THR500 | ASP355 | 66 |
| | 4 | GLY496 | LYS353 | 64 |
| | 5 | LYS417 | ASP30 | 57 |
| | 6 | GLN493 | LYS31 | 56 |
| | 7 | TYR505 | GLU37 | 50 |
| | 8 | ASN487 | TYR83 | 41 |
| Y453A | 1 | GLY502 | LYS353 | 84 |
| | 2 | ASN487 | TYR83 | 78 |
| | 3 | GLN493 | GLU35 | 77 |
| | 4 | TYR505 | GLU37 | 67 |
| | 5 | THR500 | ASP355 | 56 |
| | 6 | TYR449 | ASP38 | 41 |
| L455A | 1 | TYR449 | ASP38 | 93 |
| | 2 | GLY502 | LYS353 | 88 |
| | 3 | ASN487 | TYR83 | 77 |
| | 4 | LYS417 | ASP30 | 71 |
| | 5 | TYR505 | GLU37 | 60 |
| | 6 | GLN493 | GLU35 | 58 |
| | 7 | GLN498 | LYS353 | 55 |
| | 8 | GLY496 | LYS353 | 40 |
| F456A | 1 | GLY502 | LYS353 | 83 |
| | 2 | ASN487 | TYR83 | 79 |
| | 3 | GLN498 | LYS353 | 75 |
| | 4 | LYS417 | ASP30 | 74 |
| | 5 | TYR505 | GLU37 | 70 |
| | 6 | THR500 | ASP355 | 69 |
| | 7 | GLN493 | GLU35 | 63 |
| Y473A | 1 | GLY502 | LYS353 | 88 |
| | 2 | ASN487 | TYR83 | 82 |
| | 3 | GLN493 | GLU35 | 77 |
| | 4 | GLN498 | ASP38 | 69 |
| | 5 | THR500 | ASP355 | 51 |
| | 6 | GLN498 | LYS353 | 46 |
| | 7 | TYR505 | GLU37 | 41 |
| A475V | 1 | GLY502 | LYS353 | 84 |
| | 2 | GLN493 | GLU35 | 83 |
| | 3 | GLN498 | ASP38 | 70 |
| | 4 | ASN487 | TYR83 | 68 |
| | 5 | THR500 | ASP355 | 64 |
| | 6 | GLN498 | LYS353 | 42 |
| G476A | 1 | GLY502 | LYS353 | 85 |
| | 2 | TYR449 | ASP38 | 78 |
| | 3 | GLN493 | GLU35 | 77 |
| | 4 | TYR505 | ALA386 | 75 |
| | 5 | GLN498 | ASP38 | 73 |
| | 6 | TYR83 | ASN487 | 70 |
| | 7 | LYS353 | GLN498 | 41 |
| G476S | 1 | GLN493 | GLU35 | 88 |
| | 2 | TYR449 | ASP38 | 84 |
| | 3 | GLY502 | LYS353 | 83 |
| | 4 | ASN487 | TYR83 | 69 |
| | 5 | THR500 | TYR41 | 55 |
| | 6 | TYR505 | GLU37 | 52 |
| | 7 | GLN498 | ASP38 | 52 |
| | 8 | GLY496 | LYS353 | 49 |
| | 9 | GLN498 | LYS353 | 46 |

**Table S3.** (continued)

| mutant | # | RBD | ACE2 | Occupancy % |
|---|---|---|---|---|
| **T478I** | 1 | TYR449 | ASP38 | 94 |
| | 2 | GLY502 | LYS353 | 89 |
| | 3 | GLN498 | ASP38 | 81 |
| | 4 | GLN493 | GLU35 | 79 |
| | 5 | ASN487 | TYR83 | 65 |
| | 6 | TYR505 | GLU37 | 63 |
| | 7 | LYS417 | ASP30 | 49 |
| | 8 | GLN498 | LYS353 | 48 |
| | 9 | THR500 | ASP355 | 42 |
| **V483A** | 1 | TYR449 | ASP38 | 94 |
| | 2 | GLY502 | LYS353 | 89 |
| | 3 | GLN498 | ASP38 | 88 |
| | 4 | GLN493 | GLU35 | 79 |
| | 5 | ASN487 | TYR83 | 74 |
| | 6 | LYS417 | ASP30 | 57 |
| | 7 | GLN498 | LYS353 | 53 |
| | 8 | TYR505 | GLU37 | 44 |
| | 9 | THR500 | TYR41 | 44 |
| **V483F** | 1 | TYR449 | ASP38 | 92 |
| | 2 | GLY502 | LYS353 | 84 |
| | 3 | ASN487 | TYR83 | 74 |
| | 4 | LYS417 | ASP30 | 65 |
| | 5 | GLN498 | ASP38 | 58 |
| | 6 | TYR505 | GLU37 | 53 |
| | 7 | GLY496 | LYS353 | 51 |
| | 8 | THR500 | TYR41 | 44 |
| | 9 | GLN498 | LYS353 | 42 |
| **E484A** | 1 | GLY502 | LYS353 | 85 |
| | 2 | GLN493 | GLU35 | 83 |
| | 3 | TYR83 | ASN487 | 72 |
| | 4 | TYR505 | GLU37 | 68 |
| | 5 | LYS417 | ASP30 | 47 |
| | 6 | THR500 | ASP355 | 46 |
| **F486A** | 1 | GLY502 | LYS353 | 82 |
| | 2 | GLN493 | GLU35 | 78 |
| | 3 | ASN487 | TYR83 | 78 |
| | 4 | THR500 | ASP355 | 74 |
| | 5 | TYR449 | ASP38 | 60 |
| | 6 | TYR505 | ALA386 | 56 |
| | 7 | LYS417 | ASP30 | 44 |
| | 8 | GLY496 | LYS353 | 41 |

| mutant | # | RBD | ACE2 | Occupancy % |
|---|---|---|---|---|
| **N487A** | 1 | THR500 | ASP355 | 87 |
| | 2 | GLY502 | LYS353 | 81 |
| | 3 | ARG403 | GLU37 | 73 |
| | 4 | GLN498 | LYS353 | 62 |
| | 5 | GLN493 | GLU35 | 61 |
| | 6 | GLU484 | LYS31 | 45 |
| **Y489A** | 1 | TYR449 | ASP38 | 96 |
| | 2 | GLY502 | LYS353 | 88 |
| | 3 | GLN493 | GLU35 | 79 |
| | 4 | THR500 | ASP355 | 68 |
| | 5 | GLN498 | ASP38 | 66 |
| | 6 | LYS417 | ASP30 | 61 |
| | 7 | TYR505 | GLU37 | 60 |
| | 8 | ASN487 | TYR83 | 53 |
| | 9 | GLN498 | LYS353 | 47 |
| | 10 | GLN493 | LYS31 | 45 |
| | 11 | GLY496 | LYS353 | 41 |
| **Q493A** | 1 | TYR449 | ASP38 | 95 |
| | 2 | GLY502 | LYS353 | 87 |
| | 3 | GLN498 | ASP38 | 80 |
| | 4 | ASN487 | TYR83 | 78 |
| | 5 | TYR505 | GLU37 | 61 |
| | 6 | GLN498 | LYS353 | 50 |
| | 7 | LYS417 | ASP30 | 49 |
| | 8 | THR500 | TYR41 | 47 |
| | 9 | GLY496 | LYS353 | 43 |
| **S494A** | 1 | GLY502 | LYS353 | 88 |
| | 2 | GLN498 | ASP38 | 87 |
| | 3 | TYR83 | ASN487 | 82 |
| | 4 | GLN493 | GLU35 | 73 |
| | 5 | TYR505 | GLU37 | 67 |
| | 6 | THR500 | ASP355 | 65 |
| | 7 | LYS417 | ASP30 | 65 |
| **S494P** | 1 | GLN493 | GLU35 | 88 |
| | 2 | TYR449 | ASP38 | 84 |
| | 3 | GLY502 | LYS353 | 83 |
| | 4 | ASN487 | TYR83 | 69 |
| | 5 | THR500 | TYR41 | 55 |
| | 6 | TYR505 | GLU37 | 52 |
| | 7 | GLN498 | ASP38 | 52 |
| | 8 | GLY496 | LYS353 | 49 |
| | 9 | GLN498 | LYS353 | 46 |

**Table S3.** (continued)

| mutant | # | RBD | ACE2 | Occupancy % |
|---|---|---|---|---|
| **G496A** | 1 | GLY502 | LYS353 | 86 |
| | 2 | GLN493 | GLU35 | 80 |
| | 3 | TYR83 | ASN487 | 72 |
| | 4 | THR500 | ASP355 | 59 |
| | 5 | TYR449 | ASP38 | 55 |
| | 6 | GLN498 | ASP38 | 44 |
| **Q498A** | 1 | TYR449 | ASP38 | 86 |
| | 2 | GLY502 | LYS353 | 84 |
| | 3 | ASN487 | TYR83 | 78 |
| | 4 | GLN493 | GLU35 | 68 |
| | 5 | LYS417 | ASP30 | 62 |
| | 6 | THR500 | ASP355 | 58 |
| | 7 | TYR453 | HIS34 | 55 |
| | 8 | TYR505 | GLU37 | 48 |
| **T500A** | 1 | GLY502 | LYS353 | 82 |
| | 2 | GLN493 | GLU35 | 75 |
| | 3 | TYR83 | ASN487 | 74 |
| | 4 | LYS353 | GLY496 | 70 |
| | 5 | GLN42 | GLN498 | 66 |
| | 6 | LYS353 | GLN498 | 61 |
| | 7 | LYS417 | ASP30 | 53 |
| | 8 | GLN498 | GLN42 | 48 |
| | 9 | TYR505 | GLU37 | 47 |
| **N501A** | 1 | TYR449 | ASP38 | 84 |
| | 2 | GLY502 | LYS353 | 80 |
| | 3 | GLN498 | ASP38 | 79 |
| | 4 | ASN487 | TYR83 | 77 |
| | 5 | GLN493 | GLU35 | 73 |
| | 6 | TYR505 | GLU37 | 63 |
| | 7 | LYS417 | ASP30 | 61 |
| | 8 | THR500 | ASP355 | 52 |
| | 9 | GLN498 | LYS353 | 44 |
| **G502A** | 1 | ASN487 | TYR83 | 77 |
| | 2 | GLN493 | GLU35 | 77 |
| | 3 | LYS417 | ASP30 | 61 |
| | 4 | TYR505 | GLU37 | 47 |
| **Y505A** | 1 | GLY502 | LYS353 | 88 |
| | 2 | ASN487 | TYR83 | 82 |
| | 3 | GLN493 | GLU35 | 78 |
| | 4 | GLN498 | ASP38 | 78 |
| | 5 | GLN498 | LYS353 | 68 |
| | 6 | THR500 | ASP355 | 67 |
| | 7 | GLN493 | LYS31 | 50 |

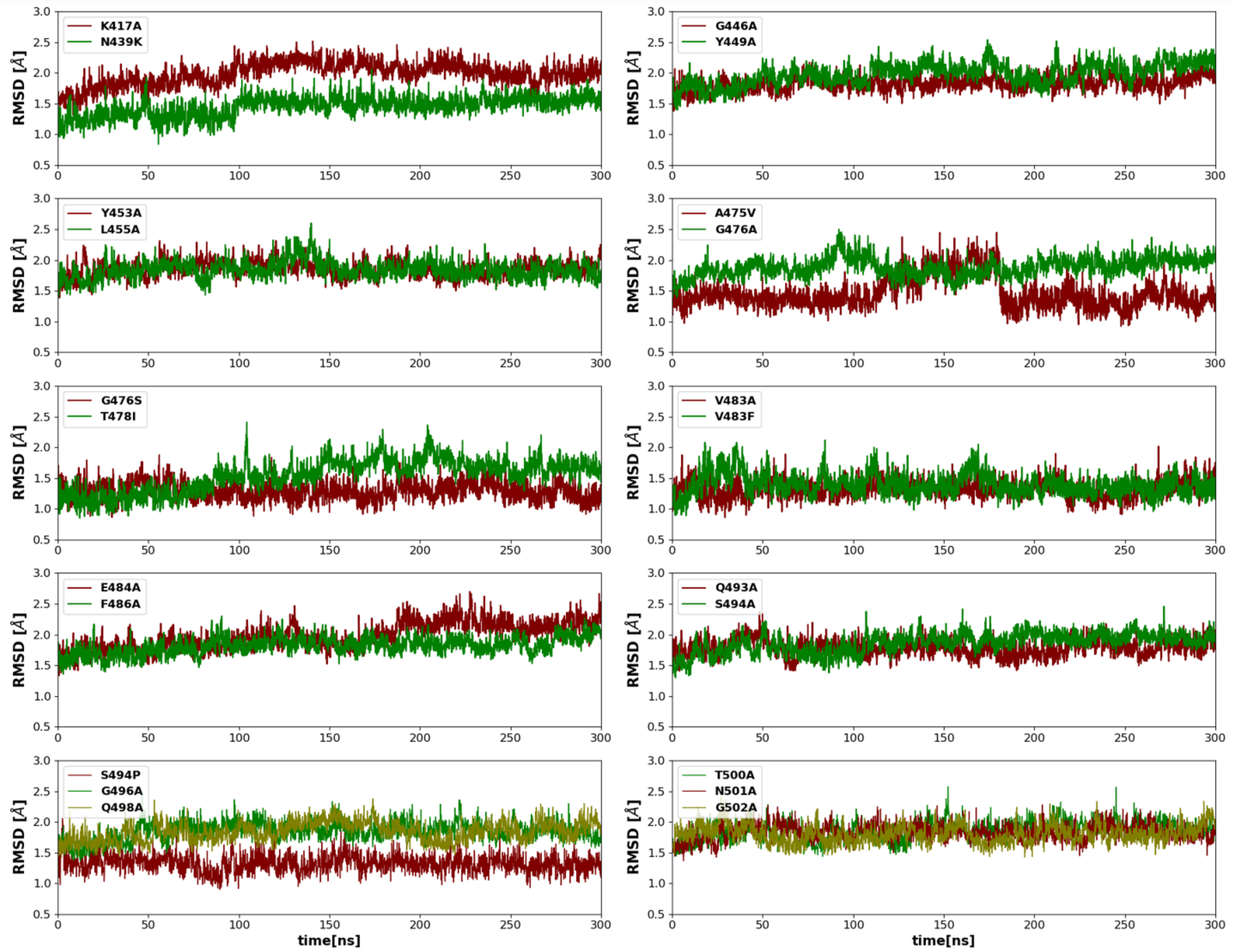


**Figure S1.** RMSD plots for mutations in nCOV-2019

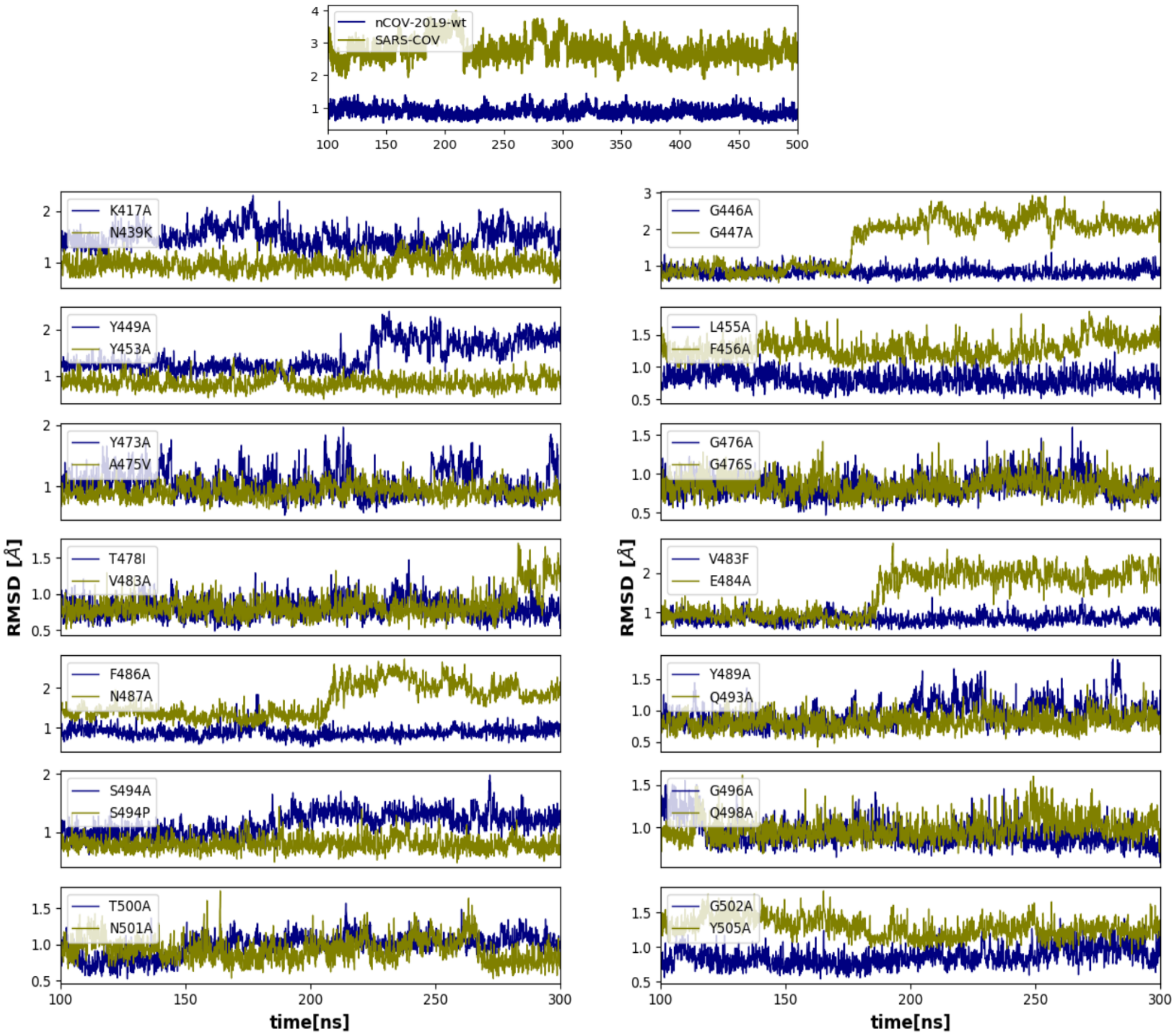


**Figure S2.** Loop RMSD for nCOV-2019, SARS-COV and all mutant systems

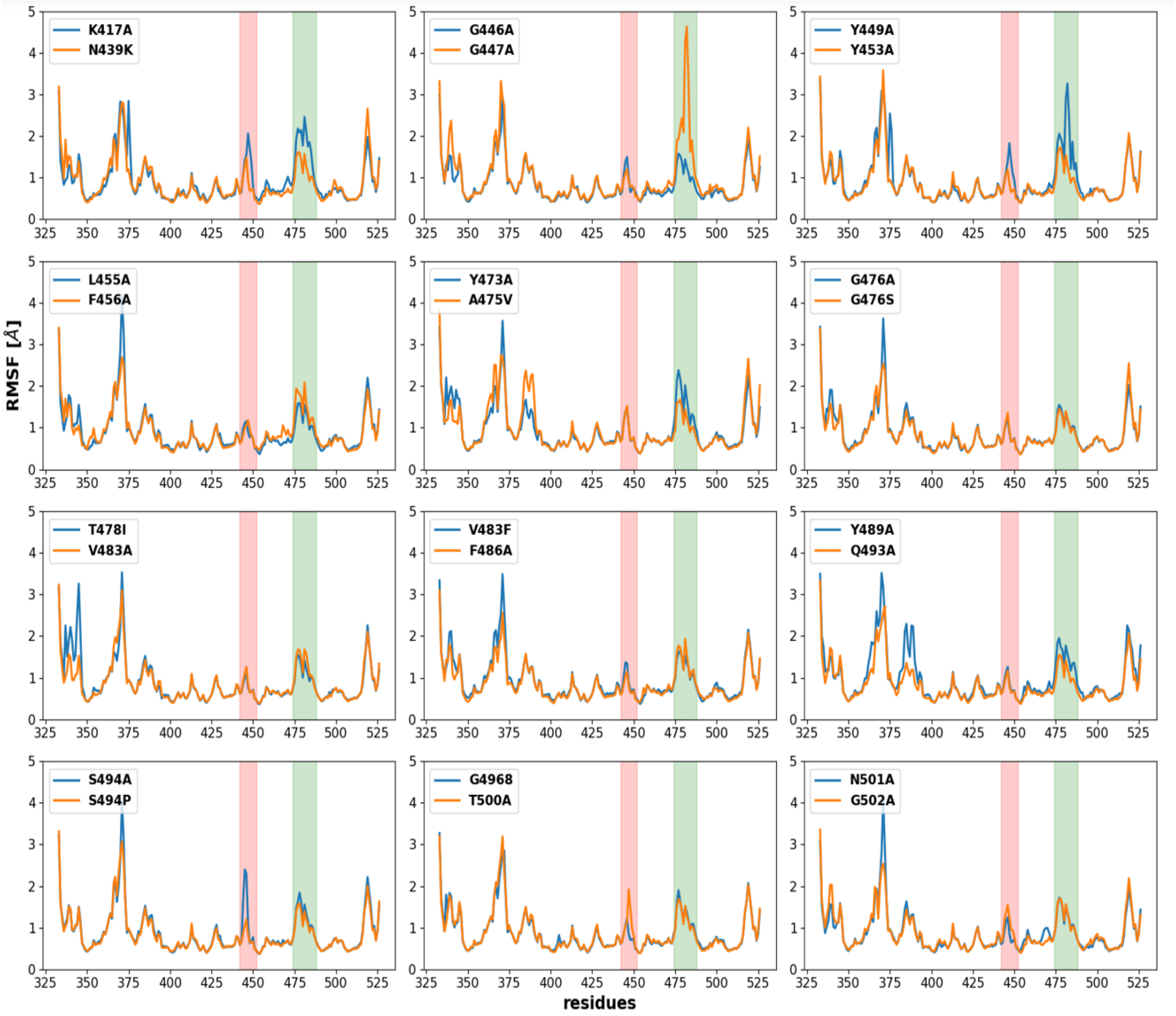


**Figure S3.** RMSF plots for mutations of nCOV-2019

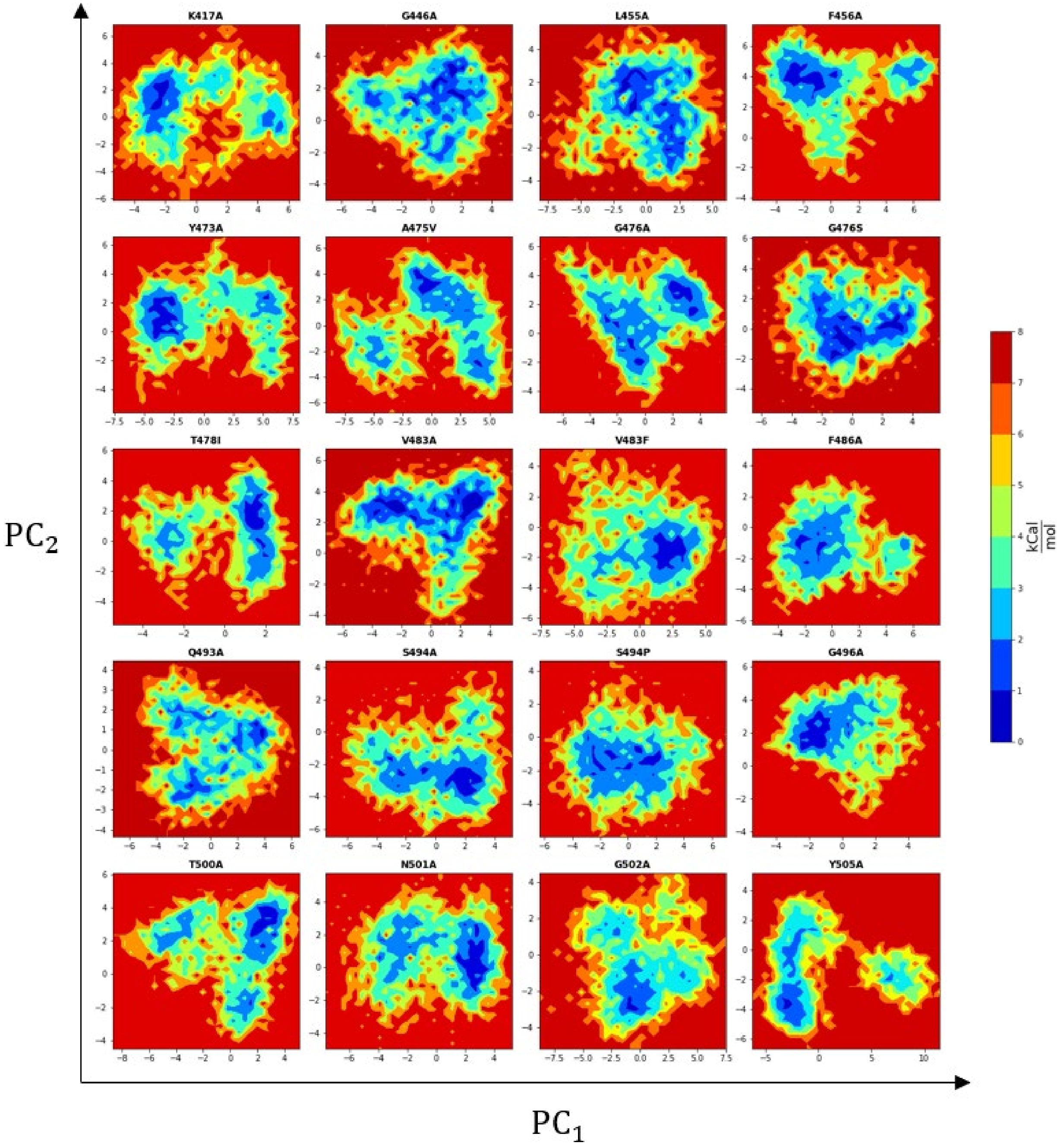


**Figure S4.** Free energy landscapes for mutants of nCOV-2019

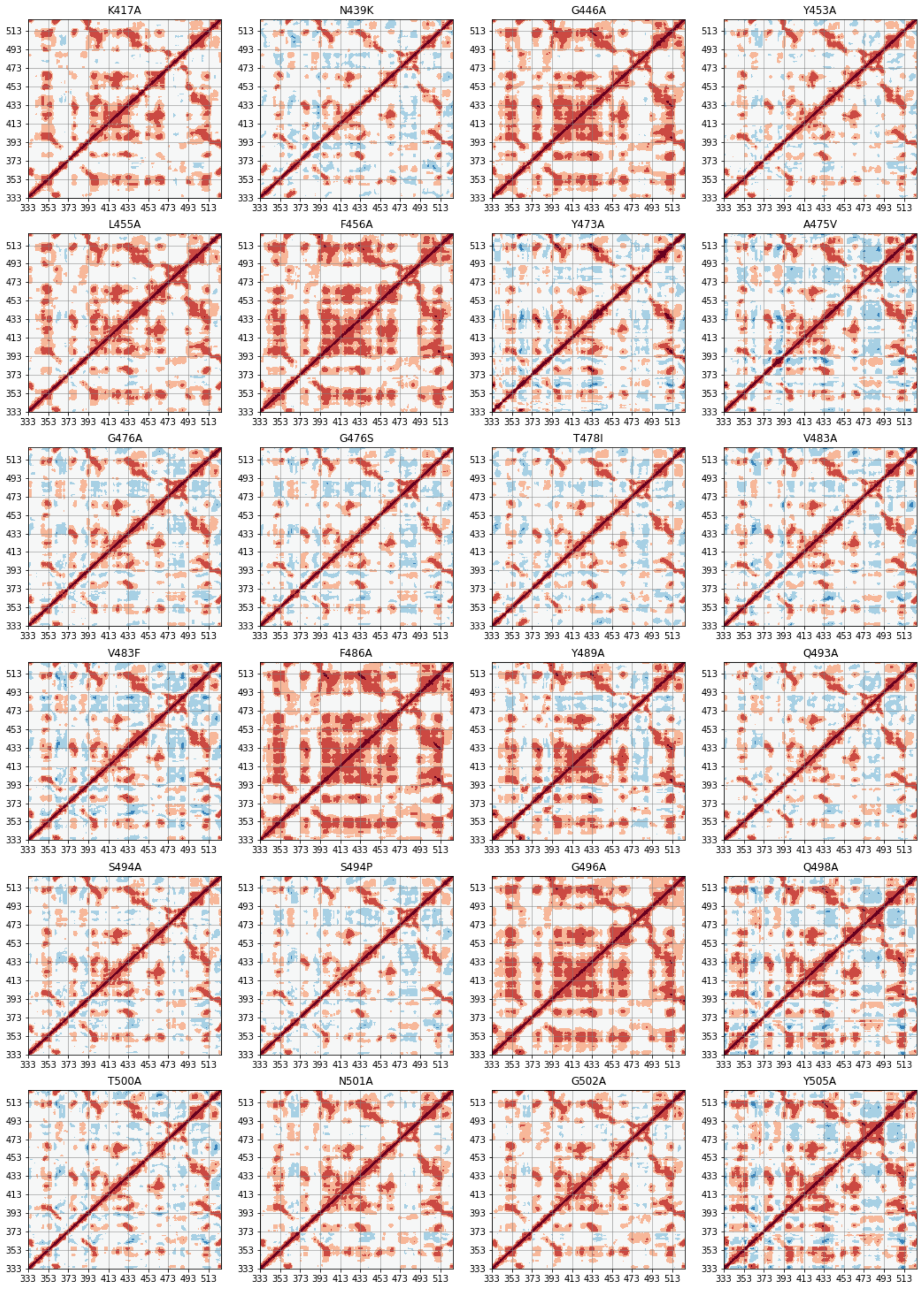


**Figure S5.** DCCM for mutants of nCOV-2019
